# Context-dependent architecture of brain state dynamics is explained by white matter connectivity and theories of network control

**DOI:** 10.1101/412429

**Authors:** Eli J. Cornblath, Arian Ashourvan, Jason Z. Kim, Richard F. Betzel, Rastko Ciric, Graham L. Baum, Xiaosong He, Kosha Ruparel, Tyler M. Moore, Ruben C. Gur, Raquel E. Gur, Russell T. Shinohara, David R. Roalf, Theodore D. Satterthwaite, Danielle S. Bassett

## Abstract

A diverse white matter network and finely tuned neuronal membrane properties allow the brain to transition seamlessly between cognitive states. However, it remains unclear how static structural connections guide the temporal progression of large-scale brain activity patterns in different cognitive states. Here, we deploy an unsupervised machine learning algorithm to define brain states as time point level activity patterns from functional magnetic resonance imaging data acquired during passive visual fixation (rest) and an n-back working memory task. We find that brain states are composed of interdigitated functional networks and exhibit context-dependent dynamics. Using diffusion-weighted imaging acquired from the same subjects, we show that structural connectivity constrains the temporal progression of brain states. We also combine tools from network control theory with geometrically conservative null models to demonstrate that brains are wired to support states of high activity in default mode areas, while requiring relatively low energy. Finally, we show that brain state dynamics change throughout development and explain working memory performance. Overall, these results elucidate the structural underpinnings of cognitively and developmentally relevant spatiotemporal brain dynamics.

## INTRODUCTION

A fundamental goal of modern neuroscience is to understand how the structure of an organism’s central nervous system influences its behavior. In the human brain, white matter fibers connect distant populations of neurons to shape the dynamic patterns of large-scale brain activity. The pattern of white matter connectivity can partially explain the spontaneous inter-regional correlations observed through resting state functional magnetic resonance imaging (fMRI)^1,2^. Network analyses of these correlations in both fMRI^3–5^ and electrophysiological data^6–8^ consistently identify groups of brain regions that display statistically similar time courses. Such groups have been variously referred to as resting state functional networks (RSNs) or putative functional modules, and coordinate similar neurological functions^5–7^.

While correlation-based approaches summarize network connectivity over a period of time, cutting-edge signal-processing approaches to fMRI can provide a richer account of brain dynamics by considering the whole-brain patterns of activity at single time points^9–15^. Such studies provide evidence for the existence of recurrent patterns of coactivation, consisting of different combinations of RSN components^9,11,12^. The study of time point level progression of coactivation patterns is intuitively complementary to the goals of brain stimulation; that is, to alleviate symptoms of neuropsychiatric illness by modulating the current and future states of specific networks^16–19^.

However, a fundamental understanding of the temporal progression of these recurrent coactivation patterns, or *brain states*, has been limited by the use of thresholding which disrupts the continuity of the time series^9–11^, a narrow focus on only a few brain regions^13^, and various modeling assumptions impacting the nature of the temporal dynamics detected^12^. Such limitations have also hampered progress in understanding how the timepoint level progression of activity patterns might be constrained by or indeed supported by underlying brain structure. Critically, the normative neurodevelopment of time-resolved brain state dynamics and their cognitive relevance also remain unknown, limiting our ability to incorporate such neurobiological features into our understanding of neuropsychiatric disorders with developmental origins^20–22^.

To address these fundamental gaps in knowledge, we consider a large, community-based sample (*n* = 879) of healthy youth from the Philadelphia Neurodevelopmental Cohort^21,23^, all of whom underwent diffusionand T1-weighted structural imaging, passive fixation resting state fMRI, and n-back working memory task fMRI^24–26^. We begin by extending unsupervised machine learning techniques to extract a set of discrete brain states from the fMRI data^10,11,14,27^, and to assign each functional volume from both rest and task scans to one of those states. We hypothesize that the temporal progression of brain states differs by cognitive task, which we test by quantifying the time that subjects dwell within states, and the propensity to transition between states. Next, we hypothesize that structural connectivity guides the temporal progression of brain states and explains why these particular brain states exist. We test these hypotheses using emerging tools from network control theory^16,28–31^, along with comparison to stringent, spatially conservative null models^32,33^. Finally, we hypothesize that brain state dynamics will change throughout development to optimize cognitive performance.

By rigorously testing these hypotheses, we show that the average time that subjects dwell in a state associated with high activity in frontoparietal cortex increases with working memory load, while the average time that subjects dwell in a state associated with high activity in default mode areas decreases. Interestingly, the probability of transitioning between brain states differed between rest and task, reflecting the ability of the brain to support context-dependent dynamics, and varied monotonically across development. Using network control theory, we show that structural brain networks are geometrically and topologically configured to specifically support the observed brain states, and not other states with similar properties. Moreover, states with many structural connections between active regions exhibit higher transition probabilities. Finally, we show that structure-function relationships and brain state dynamics change throughout development and explain individual differences in working memory performance. Overall, we demonstrate the utility of frame-level analyses in understanding the structural basis for developmentally and cognitively relevant context-dependent brain dynamics.

## METHODS

### Participants

Resting state functional magnetic resonance imaging (fMRI), n-back task fMRI, and diffusion tensor imaging (DTI) data were obtained from *n* = 1601 youth who participated in a large community-based study of brain development, known as the Philadelphia Neurodevelopmental Cohort (PNC)^34^. Here we study a sample of *n* = 879 participants between the ages of 8 and 22 years (mean = 15.9, s.d. = 3.3,386 males, 493 females) with high quality diffusion imaging, rest BOLD fMRI, and n-back task BOLD fMRI data. Our sample only contained subjects with low estimated head motion and without any radiological abnormalities or medical problems that might impact brain function (see Supplementary Methods for detailed exclusion criteria).

### Functional Scan Types

During the resting-state scan, a fixation cross was displayed as images were acquired. Subjects were instructed to stay awake, keep their eyes open, fixate on the displayed crosshair, and remain still. Total resting state scan duration was 6.2 minutes. As previously described^35^, we used the fractal n-back task^36^ to measure working memory function. The task was chosen because it is a reliable probe of the executive system and is not contaminated by lexical processing abilities that also evolve during adolescence^37,38^. The task involved presentation of complex geometric figures (fractals) for 500 ms, followed by a fixed interstimulus interval of 2500 ms. This occurred under the following three conditions: 0-back, 1-back, and 2-back, producing different levels of working memory load. In the 0-back condition, participants responded with a button press to a specified target fractal. For the 1-back condition, participants responded if the current fractal was identical to the previous one; in the 2-back condition, participants responded if the current fractal was identical to the item presented two trials previously. Each condition consisted of a 20-trial block (60 s); each level was repeated over three blocks. The target/foil ratio was 1:3 in all blocks, with 45 targets and 135 foils overall. Visual instructions (9 s) preceded each block, informing the participant of the upcoming condition. The task included a total of 72 s of rest, while a fixation crosshair was displayed, which was distributed equally in three blocks of 24 s at the beginning, middle, and end of the task. Total task duration was 693 s. To assess performance on the task, we used *d*′, a composite measure that takes into account both correct responses and false positives to separate performance from response bias^39^.

### Imaging data acquisition and preprocessing

MRI data were acquired on a 3 Tesla Siemens Tim Trio whole-body scanner and 32-channel head coil at the Hospital of the University of Pennsylvania. High-resolution T1-weighted images were acquired for each subject. For diffusion tensor imaging (DTI), 64 independent diffusion-weighted directions with a total of 7 *b* = 0 s/mm^2^ acquisitions were obtained over two scanning sessions to enhance reliability in structural connectivity estimates^23^. All subjects underwent functional imaging (TR = 3000 ms; TE = 32 ms; flip angle = 90 degrees; FOV = 192 *×* 192 mm; matrix = 64 *×* 64; slices = 46; slice thickness = 3 mm; slice gap = 0 mm; effective voxel resolution = 3.0 *×* 3.0 *×* 3.0 mm) during the resting-state sequence and the n-back task sequence^23,40^. During resting-state and n-back task imaging sequences, subjects’ heads were stabilized in the head coil using one foam pad over each ear and a third pad over the top of the head in order to minimize motion. Prior to any image acquisition, subjects were acclimated to the MRI environment via a mock scanning session in a decommissioned scanner. Mock scanning was accompanied by acoustic recordings of gradient coil noise produced by each scanning pulse sequence. During these sessions, feedback regarding head motion was provided using the MoTrack motion tracking system (Psychology Software Tools, Inc., Sharpsburg, PA).

Raw resting-state and n-back task fMRI BOLD data were preprocessed following the most stringent of current standards^24,41^ using XCP engine^42^: (1) distortion correction using FSL’s FUGUE utility, (2) removal of the first 4 volumes of each acquisition, (3) template registration using MCFLIRT^43^, (4) de-spiking using AFNI’s 3DDESPIKE utility, (5) demeaning to remove linear or quadratic trends, (6) boundary-based registration to individual high-resolution structural image, (7) 36parameter global confound regression including framewise motion estimates and signal from white matter and cerebrospinal fluid, and (8) first-order Butterworth filtering to retain signal in the 0.01 to 0.08 Hz range. Following these preprocessing steps, we parcellated the voxel-level data using the 463-node Lausanne atlas^44^. We excluded the brainstem, leaving 462 parcels. Our choice of parcellation scale was motivated by prior work showing that parcellations of this scale replicate voxelwise clustering results more than coarser scales with fewer parcels^10^. We excluded any subject with mean relative framewise displacement > 0.5 mm or maximum displacement > 6 mm during the n-back scan, and mean relative framewise displacement > 0.2 mm for the resting state scan.

All DTI datasets were subject to a rigorous manual quality assessment protocol that has been previously described^25^. The skull was removed by applying a mask registered to a standard fractional anisotropy map (FMRIB58) to each subject’s DTI image using an affine transformation. The FSL EDDY tool was used to correct for eddy currents and subject motion and rotate diffusion gradient vectors accordingly. Distortion correction was applied using FSL’s FUGUE utility. DSI studio was then used to estimate the diffusion tensor and perform deterministic whole-brain fiber tracking with a modified FACT algorithm that used exactly 1,000,000 streamlines per subject excluding streamlines with length < 10 mm^28,45^. Lausanne 463-node atlas parcels were extended into white matter with a 4 mm dilation^28,45^ and then registered to the first *b* = 0 volume using an affine transform^25,45^. For all analyses, the interregional streamline count between parcels divided by the mean of the parcel volumes^46^ was used as our measure of structural connectivity.

### Unsupervised clustering of BOLD volumes

In order to characterize the temporal progression of recurrent spatial patterns of high-amplitude brain activity, we began by concatenating all functional volumes into one large data matrix. Specifically, we took all brainwide patterns of BOLD activity from the resting-state scan and from the n-back task scan from all subjects, and we placed them into a matrix **X** with *N* observations (rows) and *P* features (columns). Here, *P* is the number of brain regions in the parcellation (462), and *N* is the number of subjects (879) *×* (120 resting state volumes + 225 n-back task volumes), summing up to *N* = 303255.

To determine canonical brain states present in these data, we performed 500 repetitions of *k*-means clustering for *k* = 2 to *k* = 18 using correlation as the algorithm’s measure of similarity^10,11,27^. To identify the optimal number of clusters, we assessed partition stability and the representation of brain states. Specifically, we aimed to choose the largest value of *k* that would provide consistent estimates of brain states that were represented at least once in every subject. We considered a partition set to be stable for a given *k* if it had a low coefficient of variation (high mean and low standard deviation) of the scaled Rand index^47^ for all combinations of partitions in the set. Because we intended to make cross-subject comparisons of state dwell times and transition probabilities as continuous measures, it was important to use partitions that identified brain states that were common across all subjects, rather than identifying many different states that were each only represented in a few subjects.

In considering the distribution of states across participants for different values of *k*, we observed that among *k* values exhibiting high partition stability (*k <* 8) (Fig. S1a-b), *k* values greater than 5 produced states that were not all represented in every subject (Fig. S1c). To maximize our sensitivity to a diverse set of states while maintaining inter-subject correspondence in state presence, we chose *k* = 5. After selecting *k* = 5, we chose to consider the partition with the lowest mean squared error of all 500 repetitions for subsequent analyses. To further validate the choice of *k* = 5, we evaluated the split reliability of the partition at this resolution (Fig. S1d-f). This analysis showed that cluster centroids and transition probability matrices were highly similar between independently clustered subject samples (see Supplementary Methods for details). Key findings are also reproduced in the Supplement for multiple values of *k*, at *k* = 5 for a second parcellation, and at *k* = 5 using cosine similarity as the algorithm’s measure of similarity.

### Analysis of spatiotemporal brain dynamics

After using *k*-means clustering to define discrete brain states, we generated names for each state based on the maximum cosine similarity to binary vectors based on an *a priori* defined 7-network partition^3^; names were generated separately for maximum cosine similarity of positive and negative state entries. These names only serve as a convenient way of referring to clusters instead of their index (i.e., 1-*k*), and have no impact on any analyses. Next, we computed subject-level state dwell times as the percentage of volumes in each scan that were classified as a particular state. We defined the transition probability between two states to be the probability of transitioning from state *i* to state *j* given that the current state is *i*. This information was encoded in a transition matrix with row sums equal to 1 and the *ij*^th^ elements of the matrix equal to the number of transitions from *i* to *j* divided by the number of occurrences of state *i*.

The state transition probability matrix houses several pieces of important information. We refer to the diagonal entries in the transition probability matrix as the persistence probabilities, because they indicate the probability of remaining in a given state, and we refer to the off-diagonal entries in the transition probability matrix as the transition probabilities, because they indicate the probability of transitioning between two distinct states. Given this structure, we were interested to determine whether the brain dynamics that we observed could occur in a uniformly random distribution of states and state transitions. To test the randomness of persistence probabilities, we generated subject-level null state time series that preserved dwell time but shuffled the order of states; for example, if the state time series was given by the vector [1 1 2 2 3 3], then we would permute the order of the entries in that vector uniformly at random. To test the randomness of transition probabilities, we generated null state time series that preserved only the states involved in transitions and reduced sequences of repeating states to a single state, so as to remove the potentially independent effects of state persistence in estimating transition probabilities; for example, if the state time series was given by the vector [1 1 2 2 3 3], then we would reduce that original vector to the new vector [1 2 3], and then subsequently permute the new vector uniformly at random.

Next, we assessed the properties of these transition matrices, such as the matrix symmetry, which reflects whether transitions from state 1 *→* 2 occur as frequently as transitions from state 2 *→* 1, and so forth for every state pair. Specifically, we quantified the asymmetry *ψ* of a *k*-by-*k* transition matrix **A** as:

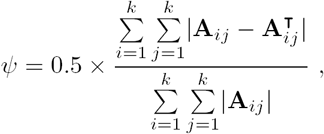

where *i* ≠ *j*. The values of this score range from 0 to 1, where 0 represents a matrix that is symmetric about the diagonal and 1 represents a matrix in which the upper triangle is –1*×* the lower triangle.

We were also interested in how much information about future states was contained within the current state. Drawing from information theoretic approaches to analysis of discrete signals, we computed the normalized mutual information (NMI) between lagged state time series to answer this question. Here, we asked whether the current state contains information about the subsequent state by computing NMI between the original state time series and a state time series lagged by one element. First, we created two copies of the state time series. Then, we removed the first element from one, **X**, and the last element from the other, **Y**, to generate two vectors of equal length such that **X** _*i*_ = **Y** _*i* +1_. We computed the NMI between **X** and **Y** as:

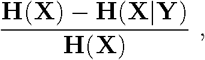

where **H**(**X**) is the entropy of **X** and **H**(**X** *|***Y**) is the conditional entropy of **X** given **Y**. Where *k* is the number of brain states,

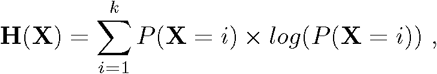

and

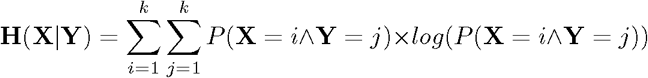

In normalizing by **H**(**X**), we ensure that the NMI ranges from 0 to 1, with 0 representing two completely independent signals and 1 representing two identical signals.

Finally, in order to assess the context-dependent nature of brain state dynamics, we performed a nonparametric permutation test to compare group-average transition probabilities between the n-back task and the resting state. First, we randomly selected two halves of the full sample. Next, we generated two group-average transition matrices by averaging together resting state transition matrices from one half and n-back transition matrices from the other half, and *vice versa*. This procedure was repeated 10000 times, and we retained the difference between the two halves at every element of the transition matrix. We generated a *p*-value for each element of the transition matrix by dividing the number of times the observed difference between n-back and rest at that element exceeded the null distribution of differences.

### Structure-function relationships

In the process of our investigation, we used several approaches to assess how white matter structural connectivity (SC) supports the complex spatiotemporal dynamics of the brain. First, we asked whether state pairs with more structural connections between their active regions also transition between one another more frequently than states pairs with fewer structural connections between their active regions. Highly active regions were defined as those exceeding a *z*-score of 1.2 in the cluster centroid, although analyses using this thresholding method are reproduced for a range of threshold values in Fig. S4. Given a weighted structural adjacency matrix **W**, we defined the *inter-state SC* as the mean value of all elements of **W** connecting highly active regions in each state, thus constructing a symmetric inter-state SC matrix **S**. Next, to assess whether state transitions occur more frequently between highly structurally connected regions, we computed the correlation between the off-diagonal elements of **S** and the off-diagonal elements of the transition probability matrix **T**. We removed the diagonal elements of both **S** and **T** because they reflect within-state SC and state persistence, respectively, rather than between-state SC and state transitions.

### Network control theory

To better understand the structural basis for the observed brain states themselves, as well as their persistence dynamics, we employed tools from network control theory^31,48^. We represent the volume-normalized, streamline-weighted structural network estimated from diffusion tractography as an *N × N* matrix **A**, where *N* is the number of brain regions in the parcellation and the elements **A**_*ij*_ contain the estimated strength of structural connectivity between region *i* and *j*, where *i* and *j* can range from 1 to *N.* Because diffusion tractography cannot estimate within-region structural connectivity, *A*_*ij*_ = 0 whenever *i* = *j*.

We allow each node to carry a real value, contained in the map *x*: 𝒩_*ℕ*0_ *←* ℝ*N*, to describe the activity at each region in each brain state. Next, we employ a linear, time-invariant model of network dynamics:

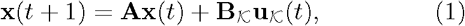

where **x** describes the state (i.e. BOLD signal) of brain regions over time, and the value of **x** _*i*_ describes the activity level of region *i*.

After stipulating this dynamical model, we computed the persistence energy **P**_*e*_ as a proxy for the stability of each clustered brain state given the white matter connections represented in **A**. Specifically, we defined **P**_*e*_ as the minimum total input energy **u**_*K*_ into every brain region required to maintain the brain state and oppose the uncontrolled trajectory away from that brain state driven by **A**. See Supplementary Methods for details on computation of minimum control energy. For the purposes of control theoretic simulations, we were interested in exploring the fundamental role of white matter architecture in supporting brain states, rather than how individual variability in white matter might impact estimates of **P**_*e*_. Thus, we constructed a single group-representative **A** generated through distance-dependent consistency thresholding^49,50^ of all subjects’ structural connectivity matrices, a process which has been described in detail elsewhere^51^.

### Spatially embedded null models

To assay the specificity of brain state activity patterns themselves, we compared **P**_*e*_ for actual brain states to **P**_*e*_ for a distribution of null brain states. We generated null states using a recent method developed to assay the specificity of regional associations in a spatially conservative manner^32^. Following this method^32^, we projected node-level data to a cortical surface, inflated the surface to a sphere using FreeSurfer tools, applied a rotation to the sphere, collapsed it back to a cortical surface, and extracted node-level data by averaging over vertices belonging to each region. This process preserves the spatial grouping and relative locations of regions with similar activity while still changing their absolute locations. Importantly, reflected versions of the same rotation are applied for each hemisphere, thus also preserving the symmetry of the original activity pattern.

To assay the specificity of our findings to real structural brain networks, we compared **P**_*e*_ estimates to a recently developed, highly spatially and topologically conservative structural network null model^33^. This model exactly preserves the degree sequence and edge weight distribution, while approximately preserving the edge length distribution and edge length-weight relationship. We also compared our findings to a more commonly used, less conservative topological null model^52^ which preserves only the degree distribution, but not degree sequence, of the network.

### Developmental and cognitive trends of brain dynamics

After identifying non-random brain dynamics at the level of individual frames, we hypothesized that features of these dynamics would change throughout normative neurodevelopment, and moreover that they would map to cognitive performance. To assess potential developmental trends of spatiotemporal brain dynamics, we fit the following model using linear regression:

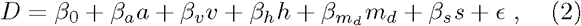

where *a* is age, *v* is total intracranial volume, *m*_*d*_ is the mean framewise displacement during rest or n-back scans, *h* is handedness, *s* is sex, *ϵ* is an error term, and *D* is a measure of brain dynamics such as dwell time, transition probability, or asymmetry. To assess potential relations between cognitive performance and spatiotemporal brain dynamics, we fit the following model using linear regression:

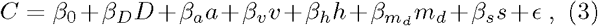

where *C* is the overall or n-back block-specific *d ′* score, which we use as our measure of working memory performance, and all other variables are the same as described above. For all analyses, we applied a Bonferroni correction for multiple comparisons, accounting for tests performed over all states or state transitions within each scan. We chose the Bonferroni-level correction because it is a conservative approach given that each state’s dwell times and transitions are not fully independent of one another.

## RESULTS

### Brain states resemble resting state functional networks

The spatiotemporal dynamics of brain activity are exceedingly complex and not fully understood. Here we used unsupervised machine learning to define clusters of statistically similar and temporally recurrent spatial activity patterns in resting and n-back task fMRI scans (Fig. 1a). Each cluster represents a canonical brain state, which we can parsimoniously study by considering the cluster centroid, or the mean activity pattern in that cluster. We found that the spatial organization of activity within the cluster centroids resembled previously reported community structure^53^ and clustered subgraphs^3,4^ in static resting-state inter-regional correlations. These resting-state networks (RSNs)^3,4^ are detected as a result of *cofluctuations* in amplitude, whether amplitude has concurrently increased, decreased, or both. For convenience, we named the five states that we observed based on their relation to these previous RSNs, and we refer to them as the DMN+, DMN-, FPN+, VIS+, and VIS-, representing high or low activity in default mode, frontoparietal, and visual networks, respectively (Fig. 2a). We also asked which RSNs were reflected in the high and low amplitude components of each brain state activity pattern separately, confirming representation of SOM (somatomotor network), DAT (dorsal attention network), and VAT (ventral attention network) (Fig. 2b).

**Fig. 1.**
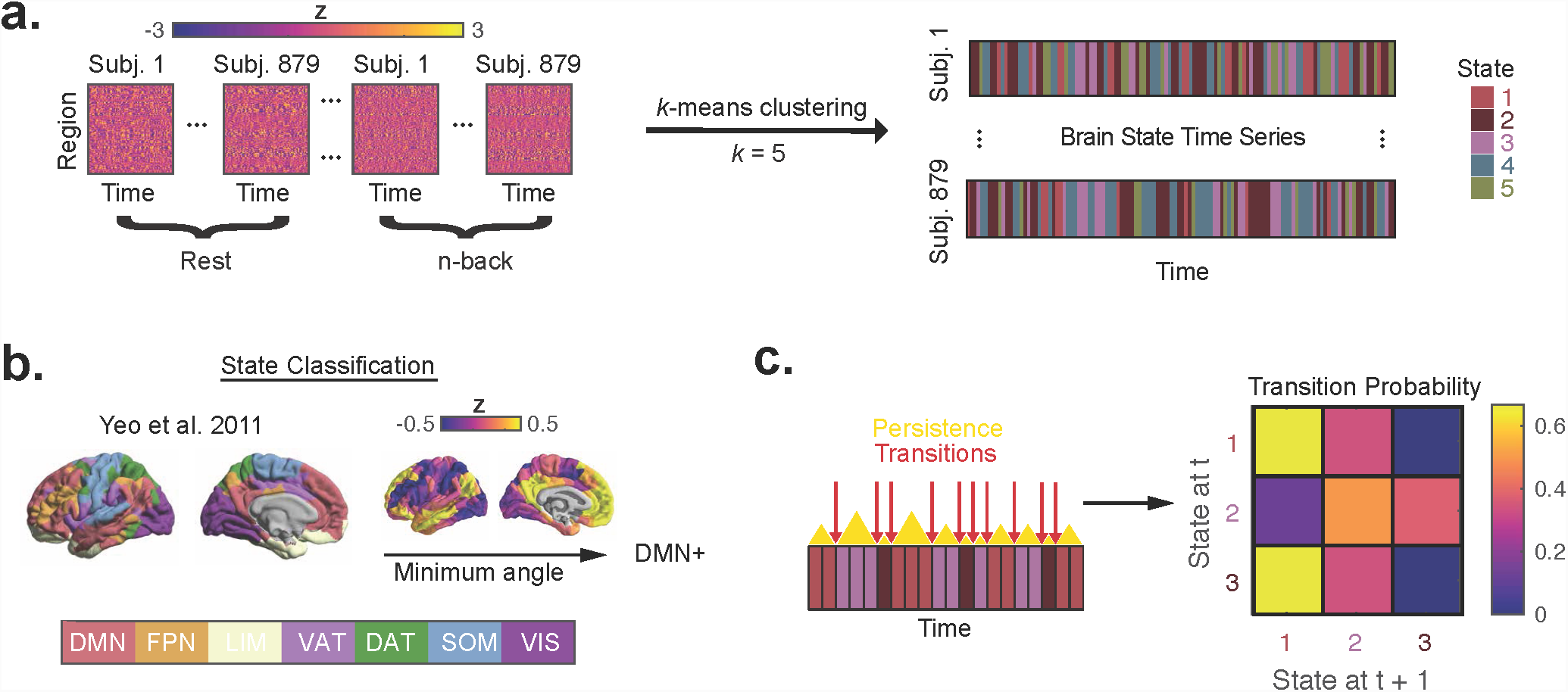
Schematic of methods for functional image analysis. *(a)* BOLD time series from resting-state and n-back task scans are concatenated across subjects. We then apply a *k*-means clustering algorithm to generate a series of cluster labels that can be mapped back to individual subjects, producing subject-specific brain state time series. *(b)* In order to characterize the spatial activity pattern within each brain state, we named each state for its maximum cosine similarity with active (+) and inactive (-) resting state networks (RSNs)^3^. *(c)* After generating labeled time series of brain states, we computed subjectspecific measures of dwell times, persistence probabilities, and transition probabilities. State transition probabilities can be represented by a matrix with rows summing to one, indicating the probability of the brain state at time *t* + 1 given the current state at time *t*. We refer to the diagonal entries of this matrix as persistence probabilities.

**Fig. 2.**
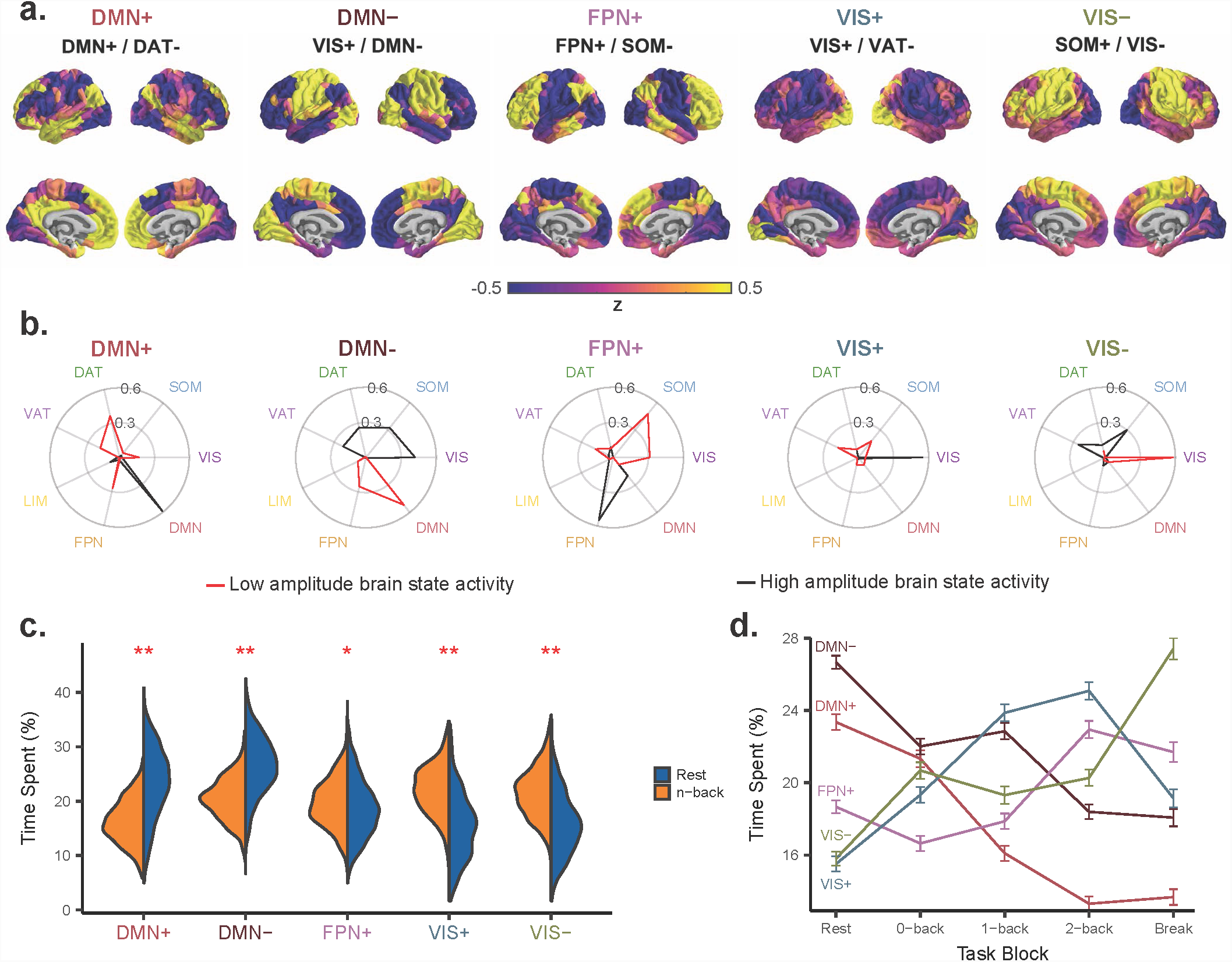
Brain states reflect community structure of resting state functional connectivity. *(a)* Brain states defined as the centroids of clusters identified using an unsupervised machine learning algorithm applied to rest and n-back task fMRI data. Brain states are labeled based on cosine similarity with *a priori* resting state functional networks^3^. The top label corresponds to the network with the most overall similarity, and the bottom two labels separated by a forward slash reflect the networks with the most similarity to the positive and negative components of each state, respectively. *(b)* Cosine similarity between positive (black) and negative (red) components of each state with binary state vectors corresponding to *a priori* definitions of functional systems^3^. Larger radial values correspond to higher cosine similarity. *(c)* Time spent, or dwell time, in each brain state differs between rest and task, with DMN states exhibiting higher dwell times during rest and VIS states exhibiting higher dwell times during task. **, *p <* 10^-15^, *, *p <* 10^-4^, Bonferroni-corrected paired *t*-tests. *(d)* Within task scans, the state dwell times change with increasing cognitive load. Interestingly, relationships between dwell time and cognitive load were state-dependent and non-monotonic. The thick colored lines indicate the mean across subjects, and the error bars represent 2 standard errors above and below the mean for each block. *DAT*, dorsal attention network, *DMN*, default mode network, *FPN*, frontoparietal network, *LIM*, limbic network, *SOM* somatomotor network, *VAT*, ventral attention network, and *VIS*, visual network.

Notably, our clustering approach captured simultaneous coactivation of multiple RSNs within each of these clustered brain states. For example, while the default mode network (DMN) exhibited high amplitude in the DMN+ state, the DAT simultaneously exhibited low amplitude in this state. These patterns of simultaneous activation and deactivation evidence rich interactions between RSNs. In addition to simultaneous coexpression of high and low amplitude RSN activity, we found that entire brain states were spatially anticorrelated with one another (Fig. S2a). Specifically, DMN+ and FPN+ were anticorrelated with DMN-, and both VIS+ and VIS- were anticorrelated with each other. These spatially anticorrelated states could represent “on” and “off” states involving similar regions^27^, potentially reflecting a singular underlying driver of oscillatory dynamics. Because of debate over whether global signal regression (GSR) can drive anticorrelated patterns in BOLD fMRI^54^, we performed the same clustering procedure on rest and n-back task BOLD time series from 100 unrelated subjects from the Human Connectome Project (HCP) without applying GSR (Fig. S3a). The brain states we identified exhibited spatial anticorrelation (Fig. S3c) and unambiguous similarity (Fig. S3b) to states identified in PNC, suggesting that GSR interacts minimally with our method for identification of brain states.

We were also interested in whether these brain states differed between rest and n-back scans. When we computed the characteristic brain state activity patterns separately for rest and n-back scans, we found that brain states were highly similar between the two conditions. The mean pairwise Pearson correlations between the two sets of states was *r* = 0.96, with a standard deviation of *r -* = 0.01 (Fig. S2b). However, the time spent in each state, or dwell time, differed between rest and n-back scans (Fig. 2c). Using paired *t*-tests, we assessed whether the population mean of subject-specific differences in brain state dwell times between n-back and rest (*µ*_*nback*_ *µ*_*rest*_) differed from 0. We observed higher dwell times in the two visual states (VIS+: *µ*_*nback*_ - *µ*_*rest*_ = 6.53, *p <* 10^-15^, VIS-: *µ*_*nback-rest*_ = 5.76, *p <* 10^-15^), and lower dwell times in the two default mode states (DMN+: *µnback-rest* = −7.09, *p <* 10*|*, DMN-: *µnback-rest* = −6.20, *p <* 10^-15^), during n-back scans.

Within n-back scans, each state’s dwell time changed differently over task blocks. Dwell times of spatially anticorrelated visual states showed opposite trends with increasing cognitive load (Fig. 2d). However, spatially anticorrelated DMN states both decreased with increasing cognitive load. As expected, the frontoparietal network (FPN) state increased from 0-back to 2-back, consistent with previous work in the same cohort that considered the activity of each voxel independently^40^. Importantly, we were able to recapitulate these results using a method based on single time point activity patterns. Collectively, these findings demonstrate that working memory load has state-dependent effects on the dwell times of common large-scale brain states, suggesting that the differential expression of brain states may be important for specific cognitive functions.

### Large-scale brain state dynamics

After demonstrating that discrete, large-scale brain states have context-dependent dwell times, we investigated the temporal dynamics of these brain states at the level of individual BOLD volumes. For each subject, we computed the probability of transitioning between each pair of brain states (Fig. 3a-b). Such transition matrices fully describe the dynamics of Markov processes^55,56^, in which the probability of states at time *t* + 1 is determined by the state at time *t*. While our method does not explicitly assume this property, we were interested in whether the brain could be described by such a process and what the properties of subject-level transition matrices could reveal about cognitively and developmentally relevant brain dynamics. Specifically, we hypothesized that the brain deviates from uniformly random dynamics to occupy states in a temporal order that facilitates cognitive functions.

**Fig. 3.**
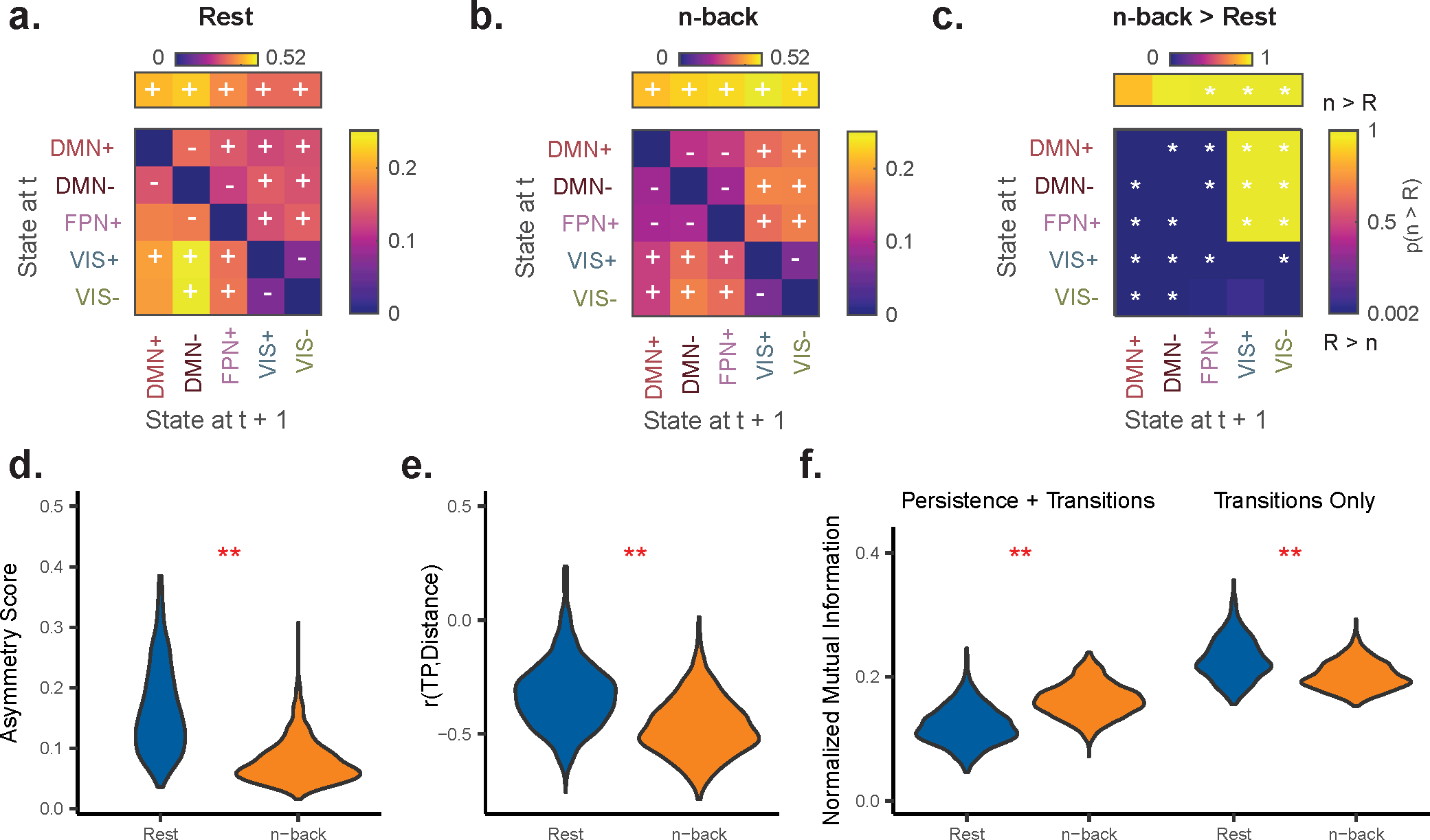
Brain state transitions are context-dependent, symmetric, and non-random. *(a-b)* Group average state transition probability matrices for rest and n-back scans. Overlayed + or indicates Bonferroni-adjusted *p <* 0.05 for transitions occurring more or less, respectively, than expected under an appropriate random null model. Persistence probabilities are removed from the diagonal and depicted above the transition matrix. *(c)* Non-parametric permutation testing demonstrating differences between the rest and n-back group average transition probability matrices. Extremes of the color axis indicate statistical significance, with larger values indicating higher transition probabilities in n-back relative to rest. *, Bonferroniadjusted *p <* 0.05 or *p >* 0.95. *(d)* Subject-level distributions of matrix asymmetry scores demonstrate that resting state transition probabilities are asymmetric relative to n-back. *(e)* Subject-level distributions of the correlation between transition probabilities and Euclidean distance between states for rest (left) and n-back (right). *(f)* Single-frame lagged, normalized mutual information for rest and n-back computed with full state time series *(left)* or transition sequence only *(right)*. **, *p <* 10^-15^. Paired *t*-tests were used in panels *(d-f)*. *TP*, transition probability.

First, we asked whether the observed persistence and transition probabilities were likely in uniformly random sequences of states and state transitions. Relative to two null models (see Methods for full details), we found that all persistence probabilities in rest and n-back occurred more than expected, while nearly all state transitions occurred more or less than expected under a transition-preserving null model (Fig. 3a-b). Interestingly, at rest and during task, VIS+ and VIS- states transitioned less frequently than expected in the random null model, suggesting that traversing between VIS+ and VISin state space involves intermediate states.

We also conducted the same analysis in an independent sample from the HCP with lower sampling rate and different preprocessing approach (see Supplementary Methods). Importantly, we found highly similar structure of transition probabilities (Rest, Pearson *r* = 0.83; n-back, Pearson *r* = 0.75; Fig. S3d-e) and persistence probabilities (Rest, Pearson *r* = 0.85; n-back, Pearson *r* = 0.84; Fig. S3d-e), suggesting that our results are generalizable and robust to preprocessing methods.

After establishing the non-random nature and robustness of brain state dynamics, we used a non-parametric permutation test to assay for differences between rest and n-back transition matrices. We found that persistence probabilities in non-DMN states were higher in the n-back scan than in the rest scan, suggesting that sequential maintenance of these activity patterns is characteristic of n-back task brain dynamics (Fig. 3c). Transitions from DMN and FPN states into VIS states were also greater during task scans than during rest scans, consistent with the higher dwell times observed in VIS states during task (Fig. 3c). These findings highlight a substantial shift in the dynamical regime that occurs between rest and a cognitively demanding task.

The ubiquitous context-dependence of brain state transitions demanded a more thorough characterization of how the brain traverses through state space at rest compared to during an n-back task. In a system with an asymmetric transition matrix, transitions from state 1 → 2 might occur much less than transitions from state 2 → 1, indicating a *directional progression* through state space. Accordingly, we assayed the overall asymmetry of each subject’s transition matrix using a normalized measure of skewness. We found that the rest transition probability matrix demonstrated more asymmetry than the n-back transition probability matrix (Fig. 3d; - *µ*_*nback-rest*_ = − 0.063, *p <* 10^-15^). This result suggests that the brain progresses through state space in a more directional fashion at rest than during the performance of the n-back working memory task.

We also hypothesized that state pairs differing less in activation magnitude would transition between one another more frequently. Consistent with our hypothesis, we found that the Euclidean distance between states was anticorrelated with the transition probability between states (Fig. 3e). Interestingly, the effect was stronger for n-back than for rest (Fig. 3e; *µ*_*nback-rest*_ = - 0.17, *p <* 10^-15^), suggesting that the brain is more prone to transition between distant states while at rest.

Finally, to further explore the non-uniformity of transitions, we measured the certainty conferred by know-ing the current brain state in predicting the next state. Specifically, we compared the lagged mutual information for brain state time series from n-back and rest. We showed that the single-frame lagged, normalized mutual information (NMI) of state time series is higher in n-back scans than in rest scans (Fig. 3f: *µ*_*nback-rest*_ = 0.041, *p <* 10^-15^), consistent with the higher persistence probabilities during n-back task performance (Fig. 3c). However, we also wished to ask the same question while controlling for the autocorrelation within the state time series due to state persistence. Therefore, we performed a secondary analysis in which we removed repeating states from the state time series (i.e. the state time series vector [1 1 2 2 3 3] becomes the new vector [1 2 3], see Methods), thereby generating a *transition sequence*. When we recomputed the single-frame lagged NMI using only the observed transition sequence, we found that rest exhibits higher lagged NMI than n-back (Fig. 3f: *µ*_*nback-rest*_ = 0.028, *p <* 10^-15^). This finding suggests that knowing the current brain state contributes more certainty in knowing the subsequent brain state at rest than during n-back.

## White matter connectivity explains brain state transition probabilities

The physical connections between neurons can in part explain their emergent function at the level of individual synapses^57^. Here we consider a much larger spatial scale and ask how the organization of white matter fibers could support the temporal progression between largescale patterns of brain activity. We hypothesized that the topology of the underlying white matter architecture would explain the observed brain state transition probabilities. Using estimates of white matter connectivity derived from diffusion-tensor imaging, we asked whether the strength of structural connectivity (SC) between highly active regions (Fig. 4a) in pairs of states (inter-state SC) was related to the transition probabilities between the respective state pair. Notably, this question is based on the notion that signals from active neuronal populations travel via white matter to effect activity changes in downstream populations. Here, we define highly active regions as those with *z*-scored activity > 1.2 in the representative brain state vector (Fig. S4c), though the results presented here hold true for a range of threshold values from (Fig. S4a-c). When we averaged transition probabilities and inter-state SC across subjects, we found that transition probabilities increased as inter-state SC increased both for the n-back task (Fig. 4b; *r* = 0.48, bootstrapped 95% confidence interval: [0.46, 0.47]) and for rest (Fig. 4c; *r* = 0.48, bootstrapped 95% confidence interval: [0.45, 0.46]).

**Fig. 4.**
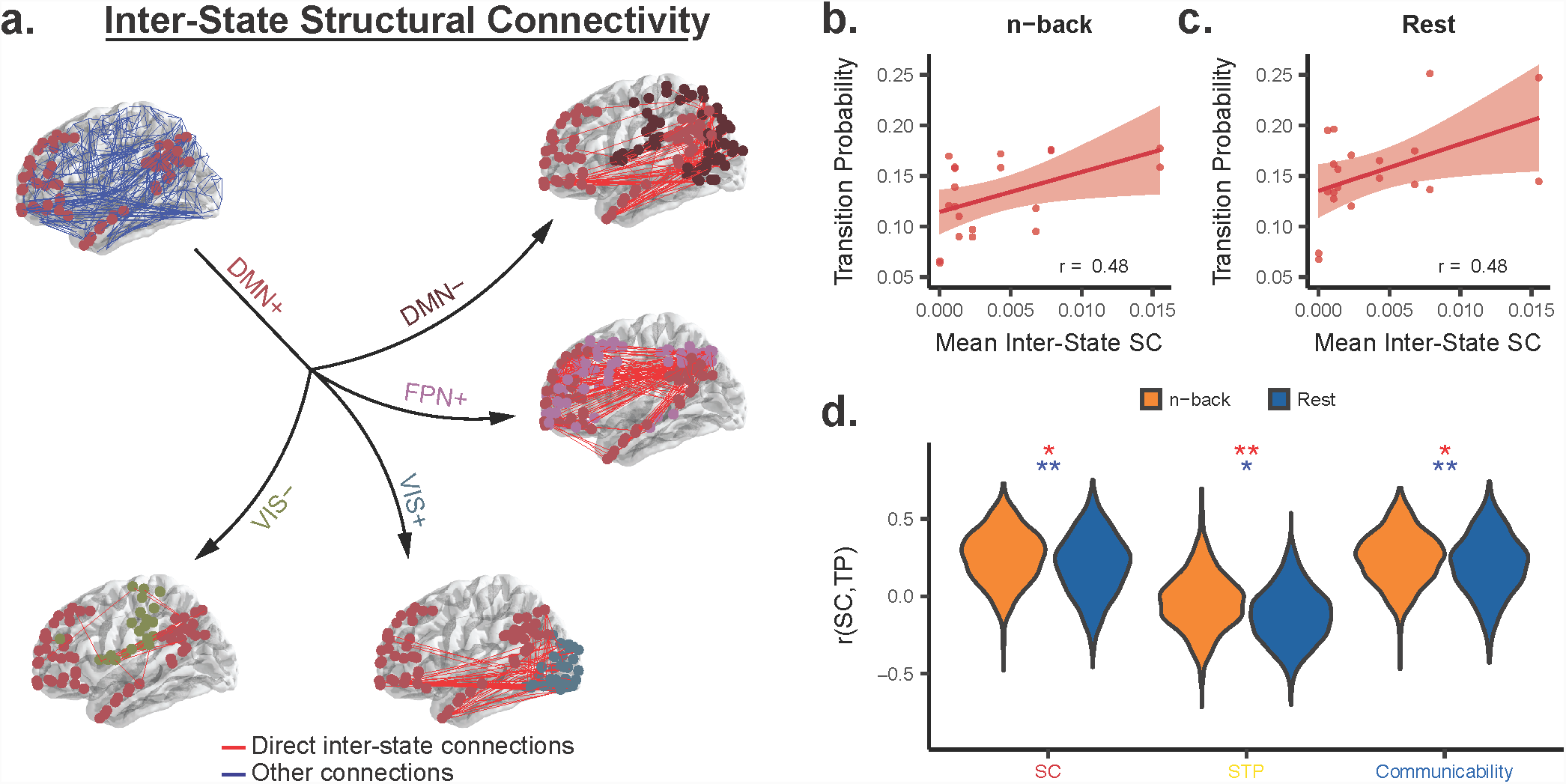
Structural connectivity between active regions predicts brain state transition probabilities. *(a)* Schematic highlighting inter-state structural connectivity (SC, red lines) between active regions in the DMN+ state (red dots) and all other states (dark red, DMN-, purple, FPN+, blue, VIS+, green, VIS-). Blue lines represent all structural connections, while red lines represent the state-specific subsets. *(b-c)* Group average inter-state SC between active regions in each pair of states is positively correlated with the group average transition probabilities for n-back *(b)* and rest *(c)* scans. *(d)* Distributions of subject-level correlations between inter-state structural metrics and transition probabilities. In all cases, subject-level correlation distributions differ from zero and are higher for n-back than rest. *TP*, transition probability. *SC*, structural connectivity. *STP*, shortest topological path. Blue *, *p <* 10^-6^ for *t*-test comparing the distribution to 0; Blue **, *p <* 10^-15^ for *t*-test comparing the distribution to 0; red *, *p <* 10^-5^ for paired *t*-test comparing rest and n-back; red **, *p <* 10^-15^ for paired *t*-test comparing rest and n-back.

Next, we computed the same correlations at the subject level using inter-state SC as well as connectivity metrics that account for indirect connectivity between regions. Evidence suggests that strong indirect connectivity between two regions can support interregional communication^58^. Specifically, we used the mean binary shortest topological path (STP)^52^ and mean communicability^59^ between highly active regions in each state. STP captures the average number of hops between two nodes, while communicability generates a weighted average of SC along walks of all lengths. We found that, for both rest and n-back, the mean subject-level Pearson correlation between the transition probabilities and each structural metric was greater than zero for SC (Fig. 4d; rest, mean *r* = 0.20, *p <* 10^-15^; n-back, mean *r* = 0.26, *p <* 10^-15^) and for communicability (Fig. 4d; rest, mean *r* = 0.20, *p <* 10^-15^; n-back, mean *r* = 0.25, *p <* 10^-15^), and was less than zero for STP (Fig. 4d; rest, mean *r* =.12, *p <* 10^-15^; n-back, mean *r* =.034, *p* = 1.8 × 10^-7^). We expected that STP would exhibit a trend opposite that of SC and communicability, because strongly connected regions often have short paths between them. Moreover, the mean subject-level correlations between inter-state SC and transition probabilities for rest (Fig. S4a) and n-back (Fig. S4b) were significantly higher than for a distribution of subject-specific, degree distribution-preserving null models^52^. Overall, these findings support the idea that state transitions are more likely to occur between states with more structural connections between their active components, the organization of which depends on higher order topological features.

We also found that these subject-level correlations were significantly higher for n-back than for rest (Fig. 4d; SC, *µ*_*nback-rest*_ = 0.055, *p* = 8.1 *×* 10^-10^; STP, *µ*_*nback-rest*_ = 0.089, *p <* 10^-15^; communicability, *× µ*_*nback-rest*_ = 0.041, *p* = 6.6 *×* 10^-6^). These findings suggest that the brain may utilize its structural connections more directly during tasks compared to during rest. Notably, our results were consistent across a range of threshold values for determining which regions were highly active (Fig. S4).

### Structural support for brain state persistence

In the previous section, we provided evidence that white matter topology guides brain state transitions. Next using techniques from network control theory, we constructed a dynamical system defined by structural connectivity and asked whether spatial and topological properties of structural brain networks specifically support the observed brain states. First, we sought to determine whether the brain states that we observed (Fig. 2a) were specifically stable in a single representative human structural brain network^49–51^ (see Methods for details). As a proxy for state stability, we computed the minimum control energy required to maintain each brain state (persistence energy) given the white matter architecture of the brain (Fig. 5c). By comparing the persistence energy for the actual states to that of null brain states^32^ with preserved symmetry and spatial clustering (Fig. 5a), we found that the minimum control energy required to maintain the DMN+ state (persistence energy) was significantly lower than for its respective null states (Fig. 5b: DMN+, *p* = 0.002, 500 sphere-permuted null states). The DMNstate also required a lower persistence energy than its respective null states, though this result was not significant after Bonferroni correction over all states (Fig. 5b, *p* = 0.012, 500 sphere-permuted null states).

**Fig. 5.**
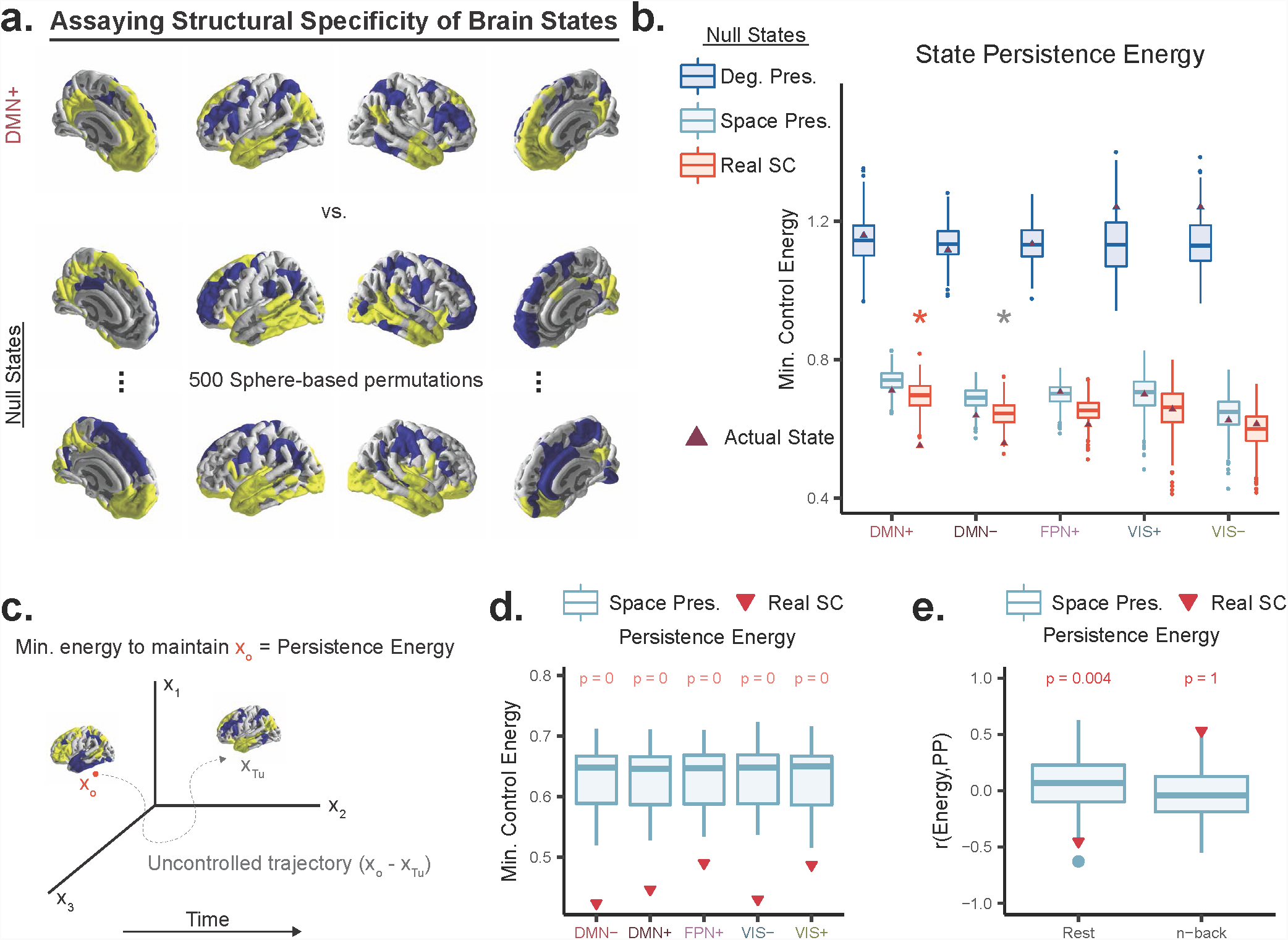
Spatial and topological properties of brain structure facilitate selective brain state stability. *(a)* Construction of null states preserving symmetry and spatial clustering of activity using sphere-based permutation^32^. We compare the minimum control energy required to maintain the brain in each state (persistence energy) relative to spatially permuted states. *(b)* We computed persistence energy for each state and its permuted variants in group average SC (orange) and found that DMN states required less energy to persist than their permuted variants. We performed the same test in two null models: a null model preserving topology (blue) and a null model preserving both topology and spatial constraints (light blue). These null models did not exhibit selective stability for actual *versus* permuted states and had overall higher minimum energies for all states. Orange *, *p <* 0.05 after Bonferroni correction. Grey *, *p <* 0.05 before Bonferroni correction. *(c)* Schematic of control theoretic simulations. A dynamical system defined by SC will follow an uncontrolled trajectory over time without any input (x_o_ - *xT*_*u*_). Persistence energy is calculated as the minimum energy input required to maintain the brain at *x*_o_ and oppose the uncontrolled trajectory. *(d)* Persistence energy required for each state in group average SC (red) or a distribution of spatially and topologically conservative null networks (light blue). For every state, the actual required persistence energy is lower in simulations predicated on the group average SC than in simulations predicated on the null model networks. *(e)* Correlation between observed persistence probabilities for rest (left) and n-back (right), and persistence energy for each state in group average SC (red) and a distribution of spatially and topologically conservative null model networks (light blue). *SC*, structural connectivity. *(Deg. Pres.)*, degree distribution-preserving null model^52^. *Space Pres.*, degree sequence, edge weight and length distribution and relationship preserving null model^33^.

We also carried out the same tests using two null models based on the group average human structural brain network: (1) a less conservative null model that preserves degree distribution only^52^ (Deg. Pres.), and (2) a highly conservative null model that preserves degree sequence, edge length distribution, edge weight distribution, and edge weight-length relationship^33^ (Space Pres.). Crucially, these null models did not exhibit selectively reduced persistence energy for observed states relative to null brain states (Fig. 5b). We also compared the persistence energy for each state in the group average SC *versus* a distribution of spatially conservative null models, and we found that group average SC had lower persistence energy for all states (Fig. 5d). Collectively, these findings suggest that the brain is geometrically and topologically configured to specifically support the observed brain states.

Next, we compared the minimum control energies of state maintenance to the observed persistence probabilities in rest and n-back. We found that persistence energies were negatively correlated with persistence probabilities at rest but positively correlated with transition probabilities during the n-back task. Importantly, the correlation between persistence energies and observed persistence probabilities was significantly stronger in real structural brain networks than in a distribution of spatially and topologically conservative null networks for both rest (Fig. 5e, *p* = 0.004, H_o_: n-back > rest) and n-back (Fig. 5e, *p* = 1, H_o_: n-back > rest). This finding suggests that the relationship between persistence energy and observed persistence probability is specific to real structural brain networks, as a result of unique geometric and topological features. Overall, these results are consistent with energetic constraints on brain state persistence at rest that are overcome during the performance of a cognitively demanding task.

### Brain state dynamics change throughout development and explain working memory performance

Developmental changes in white matter, grey matter, functional networks, and task-related activations accompany changes in behavior and cognition^40,60–63^. However, it is unclear how time point level dynamics of activity patterns and their supporting structural features contribute to these cognitive and behavioral changes. Given that the spatiotemporal brain dynamics identified by our approach have clear structural underpinnings, we hypothesized that these dynamics would change throughout normative neurodevelopment in support of emerging cognitive abilities^64^.

We used multiple linear regression to ask whether age predicted state dwell times in a context-dependent fashion while controlling for brain volume, handedness, head motion, and sex as potential confounders. Interestingly, we found that dwell times in FPN+ and DMN+ states exhibited context-dependent developmental trends (Fig. 6a). FPN+ dwell time increased with age for the nback task only (Fig. 6a; standardized *β*_*age*_ = 0.17, *p* = *×* 9.9 10^-7^), and DMN+ dwell time increased with age for rest only (Fig. 6a; standardized *β*_*age*_ = 0.12, *p* = 4.5 *×* 10^-4^).

**Fig. 6.**
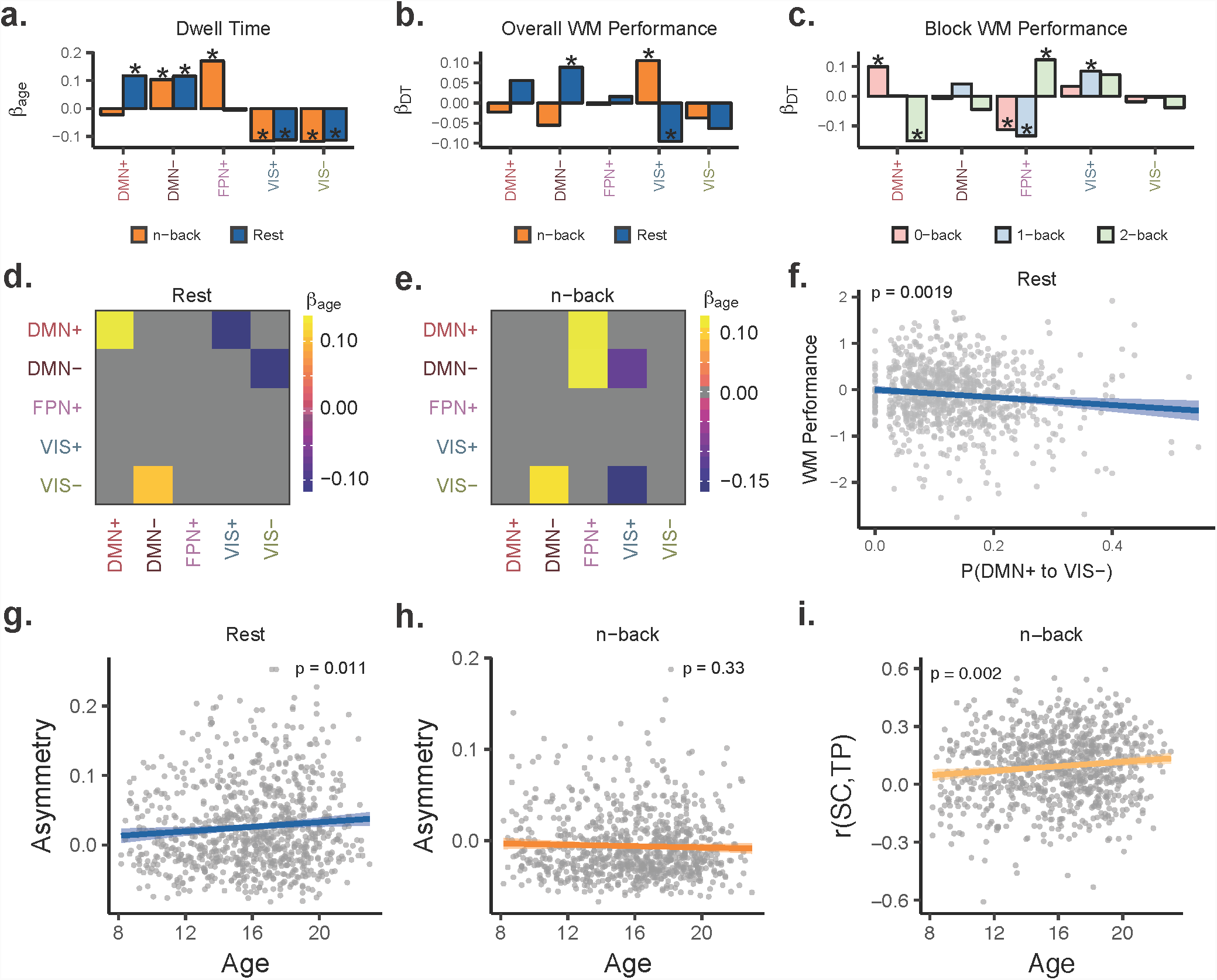
Brain state dynamics change throughout development and map to cognition. *(a)* Standardized linear regression *β* weights for age as a predictor of dwell time in each state during rest and during n-back task performance. *(b)* Standardized linear regression *β* weights for state-specific dwell time on overall working memory (WM) performance during both rest portions and non-rest portions of the n-back task scans. *(c)* Standardized linear regression *β* weights for state-specific dwell time on WM performance for each task block requiring an increasing WM load (0-back, 1-back, and 2-back). We found opposing trends for DMN+ and FPN+ states from 0-back to 2-back. *(d-e)* Standardized linear regression *β* weights for transition probabilities on age for rest (panel *(d)*) and n-back (panel *(e)*). Developmental trends of state transition probabilities differ between rest and n-back. Colored values on heatmap indicate *p <* 0.05 after Bonferroni correction. *(f)* Regression of overall WM performance on brain state transition probabilities. Only trends that were significant after Bonferroni correction are shown here; namely, transitions from DMN+ to VISat rest were negatively associated with overall WM performance. *(g-h)* Regression of asymmetry scores on age for rest (panel *(g)*) and n-back (panel *(h)*). *(i)* Subject-level correlations between inter-state SC and n-back transition probabilities regressed on age while controlling for brain volume, handedness, head motion, and sex. As age increased, the correlation between inter-state SC and n-back state transitions also increased. Regressions on cognitive performance in *(b-c,f)* controlled for age, brain volume, handedness, head motion, and sex. Points in *(f-h)* indicate subject partial residuals with respect to predictor variable on *x -*axis. *(a-c)* *, *p <* 0.05 after Bonferroni correction. *DT*, dwell time. *WM*, working memory. *SC*, structural connectivity. *TP*, transition probability.

We also assessed the developmental trends of transition probabilities. Using multiple linear regression, we tested whether age predicted each transition or persistence probability in rest and n-back, while controlling for brain volume, handedness, head motion, and sex. Similar to the context-dependent developmental trends that we observed with dwell times, we found different sets of developmental trends in rest and n-back (Fig. 6d- e), encompassing both increases and decreases in state transition probabilities. The probability of transitions from both DMN(Fig. 6e; standardized *β*_*age*_ = 0.13, *p* = 2.1 *×* 10^-4^) and DMN+ (Fig. 6e; standardized *β*_*age*_ = 0.13, *p* = 1.8 *×* 10^-4^) into FPN+ during the nback task increased with age. This observation is particularly interesting in light of prior work implicating DMN and FPN in working memory performance^40,65^. The result also provides critical evidence for the importance of direct switching between DMN and FPN states. The presence of asymmetric, context-dependent developmental trends of state transition probabilities is consistent with two distinct dynamical regimes whose energy land-scapes change independently during development. Key differences in these regimes involve the DMN and FPN, sets of regions whose activity appears to become optimized for cognition throughout development^40,66,67^.

Because DMN suppression and FPN activation have been heavily implicated in working memory^65^, we hypothesized that brain state dynamics involving the DMN and FPN would predict working memory performance. We found that increasing DMN- dwell time at rest positively predicted overall working memory performance (Fig. 6b; standardized *β*_*DT*_ = 0.089, *p* = 6.4 *×* 10^-3^). This finding might indicate that occupancy of DMN-suppressed states at rest is a signature of brain dynamics that is favorable for working memory function. Similarly, DMN+ to VIS- transitions at rest were negatively associated with overall working memory performance (Fig. 6f; standardized *β*_*TP*_ = − 0.10, *p* = 1.9 × 10^-3^), which may represent a trajectory through state space that is unfavorable for working memory function.

Surprisingly, we did not find associations between FPN+ or DMN+ activity during task and working memory performance. One possible explanation for this null finding is the presence of changes in cognitive demands between each task block^40^. To determine whether evidence supported this explanation, we examined associations between block-specific state dwell times and block-specific working memory performance (Fig. 2c-d). As expected, we found that increasing FPN+ dwell time (Fig. 6c; standardized *β*_*DT*_ = 0.12, *p* = 1.2 *×* 10^-4^) and decreasing DMN+ dwell time (Fig. 6c; standardized *β*_*DT*_ = −0.15, *p* = 2.1 *×* 10^-6^) were associated with working memory performance during the 2-back block. However, for 0-back blocks, these trends were reversed (Fig. 6c; 0-back FPN+,standardized *β*_*DT*_ = − 0.11, *p* = 6.6 *×* 10^-4^; 0-back DMN+,standardized *β*_*DT*_ = 0.10, *p* = 3.2 *×* 10^-3^). This pattern of results might reflect the engagement of alternative systems for low difficulty tasks by strong performers, thus introducing a layer of complexity to the notion of DMN and FPN as primary task-negative and task-positive systems^68^.

Finally, we tested for developmental trends of transition matrix properties (asymmetry) and structurefunction relationships. Because the developmental trends of individual transition probabilities exhibited context-dependence and asymmetry, we hypothesized that the overall transition matrix asymmetry might increase with age, reflecting the emergence of directional trajectories in state space that support cognition. We found that the asymmetry of transition probabilities at rest increases with age (Fig. 6g; standardized *β*_*age*_ = 0.086, *p* = 0.011), while the asymmetry of transition probabilities during the n-back task did not change with age (Fig. 6h; standardized *β*_*age*_ = - 0.035, *p* = 0.33). We tested for developmental trends in the relationship between inter-state SC and brain state transitions for each subject and found that for n-back only, the correlation between inter-state SC and transition probabilities increased with age (Fig. 6i; standardized *β*_*age*_ = 0.11, all *p* = 0.002). This finding suggests that context-dependent increases in structurefunction coupling accompany normative neurodevelopment, possibly reflecting the development of structural motifs that support working memory^31^.

## DISCUSSION

In the present study, we examine context-dependent, developmental trends in time point level progression of whole brain activity patterns in individuals, and demonstrate a structural basis for large-scale brain activity patterns and their dynamic temporal evolution. Using a diverse array of techniques from machine learning, network neuroscience, stochastic processes, and network control theory, we generated new insights into the complex relationship between brain structure, spatiotemporal patterns of brain activity, and behavior.

### Structural support for brain state dynamics

Cognitive functioning relies on interactions between distributed brain networks. These networks emerge as communities from the inter-regional correlations in fMRI^3–5^ and electrophysiological data^6–8^. In this paper, we identified brain states at single time points comprised of combinations of active and inactive brain networks, and we described the non-random progression of these states in time. This work adds to a body of literature suggesting that coactivation of brain networks at relatively short temporal scales evidences rich functional interactions supporting behavior^9,12,14,27^. For instance, we showed that the DMN+ state contains reduced activity in the dorsal attention network (DAT), consistent with time-resolved^9^ and correlation-based analyses^69,70^ of the DMN and DAT. The DMN+ state was present more frequently during rest than n-back, and its dwell time decreased with increasing cognitive load, consistent with the DMN’s putative role as a task-negative network^65,71^. Nevertheless, one of our major contributions to this literature lies in linking brain states themselves and their progression through time to the spatial and topological properties^33,52^ of white matter tracts.

Previous applications of network control theory have linked structural connectivity to the state space of fMRI activity patterns^72^. We expanded upon this work by showing that DMN brain states were specifically stable given the structural connectivity of the brain relative to null brain states with similar properties^32^. Moreover, the differences in observed state persistence probabilities between DMN states and other states were explained by structure-based predictions of the energy required to maintain each state. Importantly, null networks preserving the spatial and topological properties of real structural brain networks required more energy to maintain the observed states and did not exhibit selective stability for those states. These findings suggest that these particular states exist as recurrent spatial activity patterns due to the unique geometric and topological configuration of white matter tracts. The finding of selective stability only in DMN states is consistent with numerous studies suggesting unique structural and functional roles for the default mode network^73,74^, disruptions of which are associated with neuropsychiatric illness^75–77^.

We also provided evidence that structural connectivity guides the temporal progression of brain states. Our hypothesis was motivated by a drive to bridge scales of study in neuroscience; at the synapse level in neuronal circuits, an increase in the firing rate of an excitatory neuron generally elicits depolarization of connected neurons^57,78^. We tested for evidence of a similar phenomenon on a larger scale by asking whether dense white matter connections between active regions in two brain states might facilitate transitions between the two states. Indeed, transition probabilities between brain states were positively correlated with the strength of direct structural connectivity between active regions in each state, both at the group and subject levels. These results are consistent with prior evidence for the importance of direct connectivity in transmitting information between brain regions^1,79^. However, our analysis differs in that we relate structural connectivity to time point level progression of large scale activity patterns measured empirically in humans.

### Context-dependent brain state dynamics

Brain state transition probabilities differed between rest and n-back, and consistent with prior reports^12^, occurred non-randomly; that is, many state transitions occurred more than or less than expected under a uniform random null model that controlled for differences in state dwell times. To gain an intuition regarding these results, it is useful to consider a representation of brain dynamics as a state space^29,30,72^ where the magnitude of local energetic minima determines state dwell time, and where the energy landscape between minima determines state transition probabilities. Our findings are consistent with the presence of biased occupancy and preferred trajectories in this state space^30,80^, and are inconsistent with the notion of traversing the space between local minima with equal probability. For instance, n-back task dynamics were more symmetric in traversing this space, with transitions mostly to nearby states, while resting state dynamics exhibited asymmetry, with transitions to distant states. Moreover, the increased certainty within the transition sequence at rest is consistent with the notion that spontaneous brain activity involves targeted exploration. These context-dependent dynamics reflect a reshaping of the whole brain state space to facilitate specific cognitive processes.

However, such a model whose energy landscape is based on static structure alone cannot fully explain the context-dependent dynamics that we observed. Changes in neuromodulatory input can alter the behavior of smaller neural circuits^81,82^, with alpha-2 adrenergic signaling specifically implicated in working memory^83,84^. Thus, changes in neuromodulatory input in response to environmental demands might trigger shifts between multiple large-scale dynamical regimes in the brain. Un-derstanding the forces that alter the dynamical regime of the brain to meet environmental demands is a fundamental goal of neuroscience. Toward this end, a similar study of temporal dynamics encompassing many different tasks would help provide a rich understanding of context-dependent energy landscapes necessary to test this hypothesis.

### Developmental changes in brain state dynamics important for cognition

Unlike previous time point level fMRI analyses^9,12,14^, our method unambiguously labels every time point in every subject for rest and n-back as belonging to a discrete, common state. We intentionally designed our method in this way to make comparisons across contexts and across subjects throughout different developmental stages. Indeed, these comparisons revealed context-specific developmental trends, suggesting that as brain structure develops, multiple trajectories through state space are supported. Our study offered insights into previously unexplored time-resolved brain dynamics in normative neurodevelopment. Neuropsychiatric illnesses such as schizophrenia, autism, epilepsy, and ADHD are increasingly considered developmental disorders, and therefore it is critical to understand the maturation of brain dynamics in healthy youth. Previous studies have shown that changes in activity and connectivity patterns in the DMN and FPN are critical for normal development of cognition^40,66,67^. Here we contribute to our understanding of these networks by demonstrating context-dependent developmental trends of DMN and FPN state dynamics (Fig. 6a,d-e). Interestingly, both dwell times and state transition probabilities exhibit context-dependent developmental trends, with DMN+ dwell times increasing with age at rest only (Fig. 6a), FPN+ dwell times increasing with age in nback only (Fig. 6a), and DMN to FPN+ transitions increasing with age during task only (Fig. 6e). Additionally, the relationship between state transitions and inter-state SC became stronger with age for n-back only, consistent with increased utilization of structural connections important for working memory by mature brains. Finally, we both reproduce^40^ and further characterize the role of DMN and FPN in working memory^65^ by showing that dwell times and transition probabilities involving DMN and FPN predict performance on the n-back task. Overall, these findings are consistent with the notion of multiple context-dependent dynamical regimes, which may mature via dissociable mechanisms.

## Methodological limitations

We acknowledge that a limitation of this study was a focus on discrete brain states with common spatial activity patterns across subjects rather than continuously fluctuating, overlapping functional modes of brain activity^9,80^. Nevertheless, this simplified approach also constituted a major strength of the study, because it allowed us to assess developmental trajectories and cognitive effects of previously unexplored brain dynamics across subjects in a large sample. Generating discrete states also allowed us to examine the brain using approaches from stochastic process theory^55,56^, including calculating transition probabilities. Importantly, our approach inherently accounts for the temporal autocorrelation within the BOLD signal^85^ by measuring the state persistence probability independently from the state transition probability.

We also intended to compare the progression of activity patterns through a common state space between rest and task^14^. Thus, we clustered activity patterns from both conditions simultaneously to ensure that we could compare transitions between common states for rest and n-back. Larger values of *k* yielded many brain states that were similar to those identified at *k* = 5, with some states predominantly existing in one scan or the other (Fig. S9). However, the relatively low sampling rate (TR = 3s) and short rest scan time (6 minutes) precluded the use of a larger number of clusters in stably estimating transition probabilities. Similarly, there likely exist meaningful differences in individual brain state topographies^86–88^ that certainly warrant further investigation. Importantly, we demonstrated that *k* = 5 yields stable cluster partitions robust to outliers (Fig. S1), and our results were consistent for multiple values of *k* (Fig. S6 and S7), a second parcellation at *k* = 5 (Fig. S8), an alternative distance metric (Fig. S5), and an independent sample with a higher sampling rate and no global signal regression (Fig. S3).

### Future directions

The novel approaches in this study pave the way for many future studies to continue to elucidate how a static structural connectome can give rise to complex, time-evolving activity patterns important for cognition. An intuitive and important application of our approach lies in the field of neurostimulation, where clinicians aim to implement targeted changes in the temporal evolution of brain activity patterns^16,19,89^ to alleviate symptoms of neuropsychiatric illness. In particular, network control theory and data-driven estimation of brain states are a powerful combination for this purpose. One could ask whether individual differences in structural connectivity explain variance in brain state dynamics, and thus response to neural stimulation. Application of these methods to electrophysiologic data would help validate the dynamics that we observed and elucidate more complex neural dynamics that are not reflected in the slow fluctuations of hemoglobin oxygenation captured by BOLD fMRI^90^. Our understanding of the structural support for context-dependent brain state organization and dynamics would be greatly enhanced by comparing rest and n-back dynamics to other cognitive tasks.

Furthermore, prior fMRI studies of healthy human brain dynamics during rest suggest that longer recording times are required for accurate estimates^88^. We would be interested in estimating brain state dwell times and transition probabilities in subjects with repeated acquisition of longer scan duration processed in native subject space to allow for individual differences in brain state topography^91^. Changes in dynamics over weeks or months might serve as a sensitive marker for concurrent changes in behavioral phenotypes^49^, enhancing our understanding of the relationship between behavior and temporally resolved brain dynamics. Finally, numerous neuropsychiatric illnesses, including schizophrenia, depression, and epilepsy, have demonstrated abnormalities in static functional connectivity^77,92^, time-varying functional connectivity^93–95^, task-related activation^96^, and structural connectivity^97,98^. We plan to use this time-resolved approach in future to investigate brain dynamics on a finer scale to understand their structural, genetic, and neurochemical underpinnings in neuropsychiatric illness.

## ACKNOWLEDGEMENTS

D.S.B. and E.J.C. acknowledge support from the John D. and Catherine T. MacArthur Foundation, the Alfred P. Sloan Foundation, the ISI Foundation, the Paul Allen Foundation, the Army Research Laboratory (W911NF-10-2-0022), the Army Research Office (Bassett-W911NF-14-1-0679, Grafton-W911NF-161-0474, DCISTW911NF-17-2-0181), the Office of Naval Research, the National Institute of Mental Health (2R01-DC-009209-11, R01 MH112847, R01-MH107235, R21-M MH-106799), the National Institute of Child Health and Human Development (1R01HD086888-01), National Institute of Neurological Disorders and Stroke (R01 NS099348), and the National Science Foundation (BCS-1441502, BCS-1430087, NSF PHY-1554488 and BCS-1631550). T.D.S. acknowledges support from the National Institute of Mental Health (R01MH107703, R01MH113550, and RFMH116920). The content is solely the responsibility of the authors and does not necessarily represent the official views of any of the funding agencies.

## SUPPLEMENTARY INFORMATION

### Sample exclusion criteria

We excluded 722 of the initial 1601 subjects for the following reasons: medical problems that may impact brain function, incidental radiologic abnormalities in brain structure, poor or incomplete FreeSurfer reconstruction of T1 images^26^, high motion in rest or n-back fMRI scans, high signal-to-noise ratio or poor coverage in task-free or n-back task BOLD images, and failure to meet a rigorous manual and automated quality assurance protocol for DTI^25^. Notably, our goal in constructing a sample was to compare structure-function relationships between contexts across all subjects in our sample. This analysis required highly stringent inclusion criteria that only included subjects with high quality data for rest BOLD, n-back task BOLD, and DTI.

### Split-halves validation of clustering

To ensure that our final clustering solution for *k* = 5 was not influenced by outliers or adversely impacted by model overfitting, we split our sample into two equal partitions 500 times and performed *k*-means clustering separately on each half of the dataset. We then matched clusters by computing the cross-correlation between both sets of centroids, and then by reordering the clusters based on the maximum correlation value for each cluster. We plotted those maximum correlation values and found that most were > 0.99, suggesting a high degree of robustness and stability in brain states (Fig. S1d). We also computed the state transition probabilities and state persistence probabilities for each half separately for rest and n-back, and then computed the correlation between the transition or persistence probabilities between the two data set partitions. Similarly, we found very high correlation values (> 0.99) for state transition probabilities and state persistence probabilities for both rest and nback (Fig S1e-f). These observations suggest that our estimates of brain dynamics are robust to outliers and consistent across different subsamples of our data.

### Impact of scan composition on brain states and dynamics

To ensure that our results were not biased by the fact that there were a larger number of n-back volumes (225 per scan) than rest volumes (120 per scan), we used the partition generated by clustering both entire scans together to compute separate centroids for volumes in rest or n-back scans. This analysis revealed a mean spatial Pearson correlation of 0.96 between corresponding centroids (Fig. S2b). Next, we generated a new sample by concatenating the first 6 minutes of the n-back task data for each subject and the entire 6 minutes of the rest data for each subject. We ran this sample through the clustering algorithm at *k* = 5 and found that the cluster centroids (Fig. S2c) were highly similar to those computed from the full sample (mean Pearson *r* = 0.99; Fig. S2d). We also computed transition probabilities using this cluster partition and identified highly similar group average transition matrix structure (rest, Pearson *r* = 0.996, nback, Pearson *r* = 0.983, Fig. S2e), suggesting that the temporal order of state labels was largely unaffected by the scan composition of the sample. Moreover, these results suggest that n-back state transitions are internally consistent.

Finally, we show that the differences between rest and n-back in the proportion of subjects with any absent states (Fig. S2f) is attenuated relative to the full sample (Fig. S1e). This suggests that in the full sample, n-back has better state representation due to better sampling, rather than poor classification of rest volumes. However, even with equal samples, rest still has more subjects with missing states, suggesting that there may be more variability in brain dynamics during rest. Importantly, there were still no subjects with absent states for rest or n-back at *k <* 5 (Fig. S2f), and there was at least 1 subject with a missing state in rest or n-back at *k >* 5 (though bars are very small in Fig. S2f). Collectively, these results support the simultaneous generation of partitions for rest and n-back volumes and the choice of *k* = 5 for analysis in the main text.

### Impact of sampling rate and global signal regression on brain states and dynamics

The BOLD data from the PNC was acquired at a sampling rate of one volume per 3 seconds^34^, which is relatively slow compared with other large data sets, including the Human Connectome Project^99^, which samples every 0.72 seconds. The standard preprocessing pipeline for this data set involves regression of head motion parameters, white matter confounds, cerebrospinal fluid confounds, and global signal from each voxel’s time series^42,100^. It is controversial whether this procedure, known as “global signal regression,” induces anticorrelation^54,101^.

Thus, we selected 100 unrelated subjects from the minimally preprocessed version of the Human Connectome Project (HCP) data set^102^ and performed the following preprocessing steps on resting state and n-back working memory task scans: (1) head motion regression, (2) linear and quadratic detrending, (3) bandpass filtering to retain the 0.01 to 0.08 Hz range, and (4) parcellation according to the 462 region Lausanne atlas. We concatenated all 405 volumes from the working memory task with the first 405 volumes from the resting state over all 100 subjects. We chose to make the number of volumes from each scan equal so that the clustering algorithm would not be biased towards one scan or the other.

Next, we performed *k*-means clustering on this matrix and computed the centroids (Fig. S3a). Every HCP centroid was maximally correlated with only one PNC centroid, and *vice versa*, allowing for unambiguous matching between the two sets of brain states (Fig. S3b). The DMN+ and DMN- states were the most similar to PNC states, while VIS+ and VIS- exhibited slightly lower correlations (Fig. S3b). DMN+ and DMN- states, as well as VIS+ and VIS- states, exhibited strong anticorrelation with each other (Fig. S3c). Nevertheless, the HCP off-diagonal elements of the transition probability matrices (i.e. transitions not persistence) for rest and n-back task were correlated with PNC transition probabilities at *r* = 0.83 and *r* = 0.75, respectively (Fig. S3d-e). Persistence probabilities for the HCP sample were correlated with PNC persistence probabilities at *r* = 0.85 for rest and *r* = 0.84 for n-back. This finding suggests that while there were differences in the spatial activity patterns of brain states, their dynamic progression through time was relatively similar. Moreover, the differences in transition probabilities between n-back and rest were highly similar in HCP and in PNC (Fig. S3e), with increased transitions from DMN and FPN states into VIS states. Overall, these findings suggest that while global signal regression and sampling rate may impact the spatial activity patterns comprising brain states to some degree, it does not impact estimation of their dynamics or the presence of spatial anticorrelation in their activity patterns.

### Calculating persistence energy using control theory

We represent the volume-normalized, streamline-weighted structural network estimated from diffusion tractography as the graph 𝒢 =(𝒱 ε), where 𝒱and εare the vertex and edge sets, respectively. Let *A*_*ij*_ be the weight associated with the edge (*i, j*), and define the weighted adjacency matrix of as *A* = [*A*_*ij*_], where *A*_*ij*_ = 0 whenever (*i, j*)*. ∉* ε We associate a real value with each node to generate a vector describing the network state, and we define the map *x*: ℕ _*≥*0_ ℝ *|*to describe the dynamics of the network state over time. Here we employ a simplified noise-free linear discrete-time and time-invariant model of such dynamics:

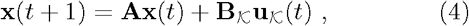

where **x** describes the state (i.e. voltage, firing rate, BOLD signal) of brain regions over time. Thus, the state vector **x** has length *N*, where *N* is the number of brain regions in the parcellation, and the value of **x** _*i*_ describes the activity level of that region. The matrix **A** is symmetric, with the diagonal elements satisfying *A* _*i i*_ = 0. Prior to calculating control energy, we divide **A** by *ξ*_0_(**A**), where *ξ*_0_(**A**) is the largest eigenvalue of **A**. We also subtract the identity matrix to prevent the system from decaying to 0 or increasing to. ∞ The input matrix **B**_*K*_ identifies the control point in the brain, where = *k*_1_*,…, k*_*m*_ and

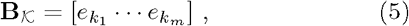

and *e* _*i*_ denotes the *i*-th canonical vector of dimension *N.* The input **u**_*K*_: ℝ _*≥*0_ ℝ *|*denotes the control strategy.

To compute the minimum control energy required to drive the system from some initial state **x**_*o*_ to some final state **x**_*f*_, we compute an invertible controllability Gramian **W**_*K*_ for the network **x**(*t* + 1) = **Ax**(*t*) + **B**_*K*_**u**_*K*_(*t*), from the set of network nodes (in our case, every node in the network), where:

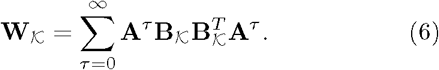

After computing the controllability Gramian, we can compute the quadratic product between the controllability Gramian and the difference between **x**_*o*_ and **x**_*f*_ to compute the minimum control energy **E_m_**:

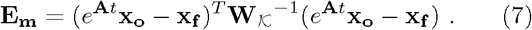

In Fig. 5, we calculate the persistence energy **P**_*e*_ as the minimum control energy where **x**_*o*_ = **x**_*f*_. We divided each cluster centroid by its Euclidean norm to control for potential effects of magnitude on persistence energy.

### Inter-state structural connectivity

We hypothesized that structural connections between highly active regions in each brain state would predict the rate of transitions between states. In order to define the highly active regions in each brain state, we computed the *z*-score of the activity in each cluster centroid, and we selected only the regions with activity greater than a threshold value in each state. We referred to these regions as *active nodes*, and determined them using the group level cluster centroids. Next, we computed the mean structural connectivity (SC) between active nodes in each pair of states for each subject, and we referred to this connectivity as the *inter-state SC*.

In Fig. 4d, we compute the correlation between subject-level inter-state SC and subject-level state transition probabilities, and we demonstrate that the distribution of these correlation values differs significantly from 0. Importantly, we excluded persistence probabilities from this analysis because (1) the large values of state persistence probabilities relative to state transition probabilities would have been outliers in these correlations, and (2) it is a separate question whether within-state connectivity predicts state persistence probability. In Fig. 4b-c, we computed the average inter-state SC and average state transition probabilities over all subjects, and we demonstrated a positive, statistically significant correlation between the two. Here, we repeat the group-average analysis carried out in Fig. 4b-c for a range of threshold values for defining active nodes, thereby demonstrating robustness to the particular choice of threshold. For both rest and n-back, the correlation between inter-state SC and state transition probability was > 0.4 for threshold values between *z* = 0.7 and *z* = 1.2 (Fig. S4a-b). Visual inspection of brain state activity patterns at different thresholds (Fig. S4c) reveals that a more stringent threshold selects very few regions, while a less stringent threshold selects the entire brain. Notably, the effect of interest was present for threshold values with activity localized to state-defining networks.

Furthermore, we tested whether the effects we saw could occur in structural networks with similar topological properties. We carried out the same analysis from Fig. S4a-b using subject-specific, degree distribution-preserving network null models^52^. Next, we used a paired *t*-test to compare the population mean SC-TP correlation between actual networks and a corresponding topological network for a range of threshold values. For both rest and n-back, we found the actual networks exhibited a higher mean correlation than the null networks (*p <* 10^-15^) for threshold values from 0.1 to 1.5 (Fig. S4d-e). This finding suggests that the relative locations of high degree nodes are important in achieving the distribution of inter-state SC values that corresponds to transition probabilities.

### Impact of cluster number, distance metric, and parcellation

A downside of *k*-means clustering is the need to specify a number of clusters, as well as a measure of distance between observations for the algorithm to minimize. We use correlation as the distance metric for all results in the body of the paper to remain consistent with prior applications of *k*-means clustering in BOLD data^10,11,14,27^. Nevertheless, we repeated our analysis at *k* = 5 using a *k*-means algorithm that maximized the cosine similarity between observations to ensure that our results were robust to the choice of distance metric. We found that cluster centroids were highly spatially similar, involving the same RSNs for each centroid (Fig. S5a). As working memory (WM) load increased during the n-back task, dwell time in DMN states decreased while FPN+ and VIS+ states increased (Fig. S5b). Transition probability matrices exhibited highly similar structure with nearly all transitions deviating from uniform random null models (Fig. S5c-d). Transitions from DMN+, DMN-, and FPN+ to VIS+ and VIS- were greater in n-back than rest (Fig. S5e), similar to (Fig. 3c). The correlation between transition probability and inter-state Euclidean distance was also more strongly negative for n-back than rest (Fig. S5f) and mean correlations between metrics of inter-state structural connectivity (SC) differed from 0 (Fig. S5g). Finally, we found highly similar associations between state dwell time, age, and WM performance. Specifically, we found that FPN+ dwell time increased with age only during n-back, while DMN+ dwell time increased with age only during rest (Fig. S5h, similar to Fig. 6a). We also found opposite trends with WM performance from 0-back to 2-back blocks for DMN+ and FPN+ states (Fig. S5i, similar to Fig. 6c).

We also reproduced our findings at different values of *k*. First, we used *k* = 4 and show centroids that capture different combinations of RSNs (Fig. S6a). The DMN+ and VIS+ states at *k* = 4 resembled DMN+ and VIS+ at *k* = 5, respectively, while SOM+ at *k* = 4 resembled DMNat *k* = 5. The VIS- state at *k* = 4 resembled FPN+ at *k* = 5, though with slightly higher activity in somatomotor regions. These findings of differential overlap are consistent with the notion of hierarchical brain states^12,14^. Accordingly, as WM load increased, we saw DMN+ and SOM+ dwell times decrease, while VISand VIS+ dwell times increased (Fig. S6b). Transitions between these states deviated from uniform randomness (Fig. S6c-d) with significant differences in transition probabilities between rest and n-back (Fig. S6e). We again observed context-dependent developmental trends at *k* = 4, with FPN+ dwell time increasing with age only during n-back and DMN+ dwell time increasing with age only during rest (Fig. S6f, similar to Fig. 6a). We also found opposite trends with WM performance from 0-back to 2-back blocks for the DMN+ state (Fig. S6i, similar to Fig. 6c). VIS-, which visually resembled FPN+ at *k* = 5, trended in a direction consistent with this phenomenon (Fig. S6g). We did not compute correlations between inter-state SC and transition probabilities at *k* = 4 because of instability in estimating correlations with only 12 observations (i.e. *k × k - k* off-diagonal elements).

At *k* = 6, cluster centroid brain states were highly similar to those at *k* = 5, with the addition of an SOM+ state and a reduction in the contribution of somatomotor regions to the DMN- state (Fig. S7a). This finding may occur due to “splitting” of the DMN- state into a somatomotor-frontoparietal subcomponent and a default mode-visual subcomponent. This appearance of “split” brain states is again consistent with hierarchical state organization^12,14^, which becomes apparent at different clustering scales. As WM load increased from 0-back to 2-back, we again saw a decrease in DMN+ and DMN-dwell times with a concurrent increase in FPN+ and VIS state dwell times (Fig. S7b). Transition probabilities at *k* = 6 deviated from uniform randomness and exhibited low transitions between VIS+ and VIS- states (Fig. S7c-d, similar to Fig. 3a-b). Moreover, transitions from DMN states into VIS states were higher in n-back than rest (Fig. S7e, similar to Fig. 3c). Similar to Fig. 3d, we found a stronger negative correlation between inter-state Euclidean distance and transition probability for rest than n-back (Fig. S7f). Using a paired *t*-test, we found that the mean correlation between 3 metrics of inter-state structural connectivity and transition probabilities in resting state and n-back significantly differed from 0 (Fig. S7g). This suggests that the relationship between inter-state structural metrics and transition probabilities is not an artifact of the choice of *k*. Finally, we found highly similar associations between state dwell time, age, and WM performance. Specifically, we found that FPN+ and SOM+ dwell times increased with age only during n-back, while DMN+ dwell time increased with age only during rest (Fig. S7h, similar to Fig. 6a). We also found opposite trends with WM performance from 0-back to 2-back blocks for DMN+ and FPN+ states (Fig. S7i, similar to Fig. 6c).

We also checked whether our findings might have been due to the choice of parcellation scale. This choice was particularly important for the relationship between transition probability and distance, where the number of dimensions in state space is explicitly determined by parcellation scale. Parcellation scale is also important for any analyses involving tractography, where relative region sizes may bias estimates of connectivity. Importantly, using the 234-node Lausanne parcellation^44^, we still found a significantly weaker negative correlation between transition probabilities and inter-state Euclidean distance for rest than n-back task (Fig. S8f). Using a paired *t*-test, we also found that the mean correlation between 3 metrics of inter-state structural connectivity and transition probabilities in resting state and n-back significantly differed from 0 (Fig. S8g). These findings suggest that parcellation scale did not bias our findings with respect to structural and distance-based constraints on transition probabilities.

---

**Fig. S1.**
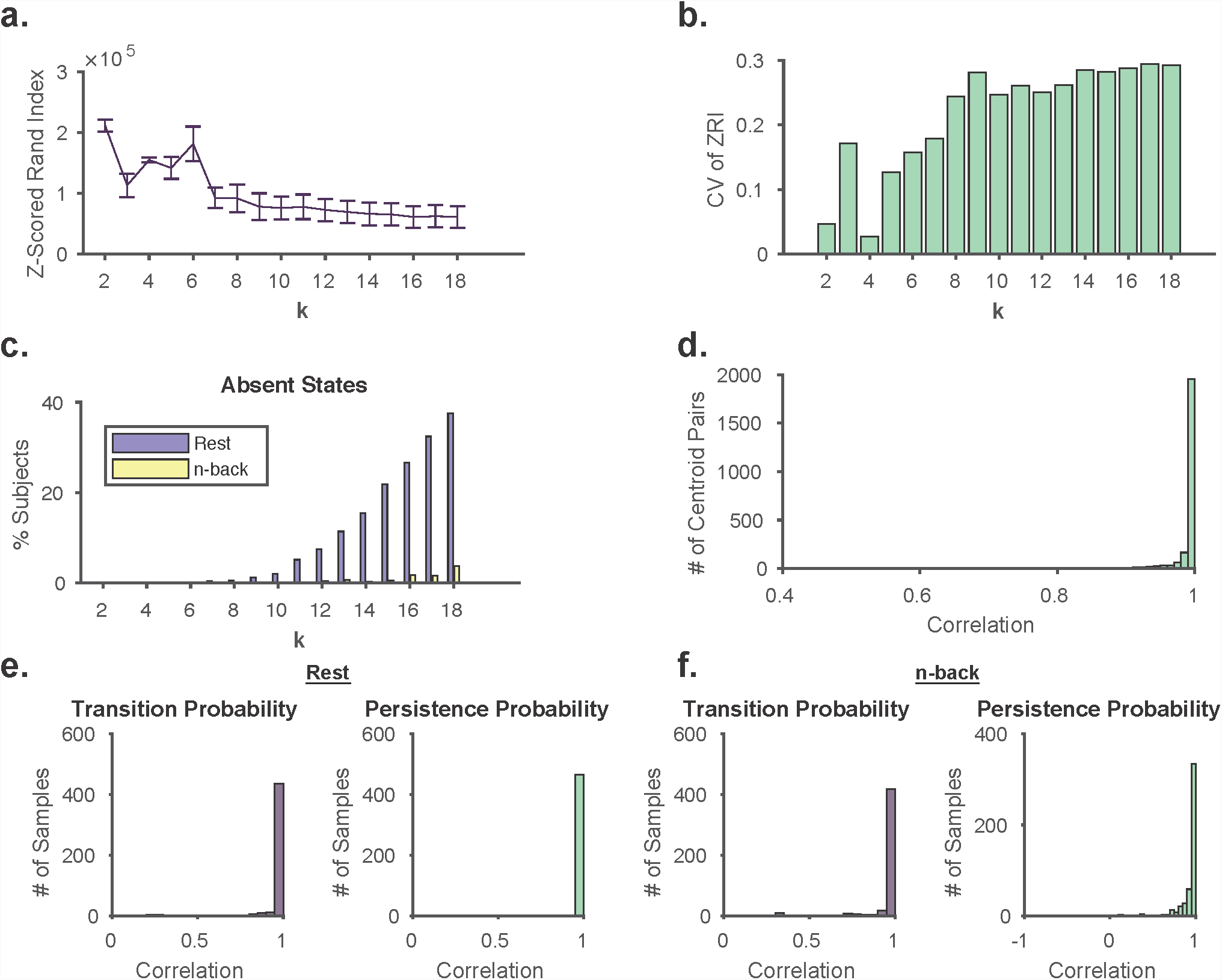
Choosing the number of clusters. *(a)* Mean (line) and standard deviation (error bars) of *z*-scored rand index (ZRI) values calculated for every pair of cluster partitions in 500 repetitions of *k*-means on concatenated rest and n-back BOLD data for *k* = 2 to *k* = 18. *(b)* Coefficient of variation (CV, given by the standard deviation divided by the mean) of the ZRI. At *k <* 8, partitions become unstable as evidenced by a high CV. *(c)* Percentage of subjects missing at least one state for rest and n-back. At *k >* 5, states begin to be incompletely represented across subjects. *(d-f)* Split-halves validation of cluster centroids *(d)*, and state transition and persistence probabilities for rest and n-back *(e-f)*. Inter-partition correlations are predominantly > 0.99, thus showing a high degree of robustness to sample composition in estimating brain states and their dynamics. *ZRI*, *z*-scored rand index. *CV*, coefficient of variation.

**Fig. S2.**
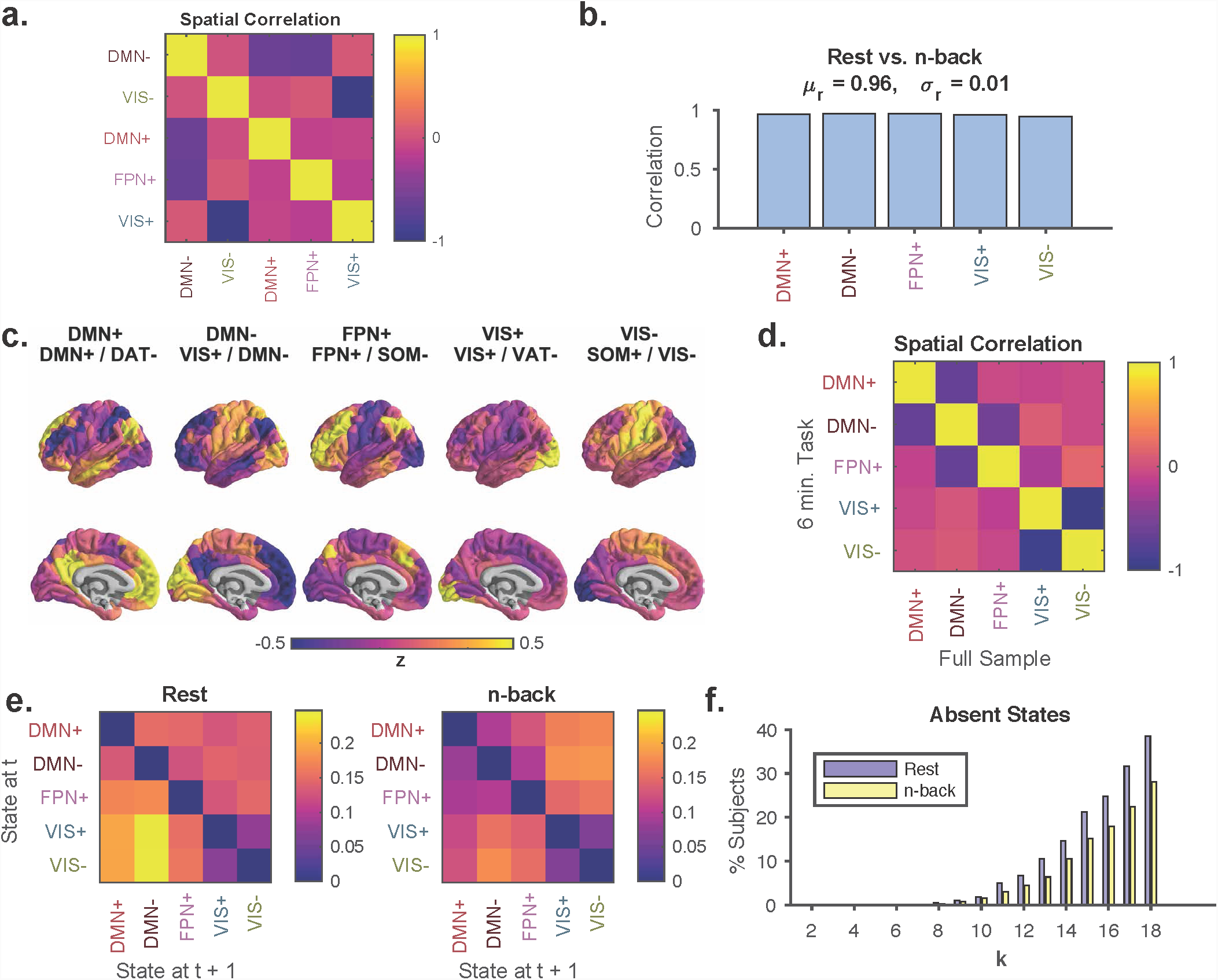
Similarity between states in rest and n-back. *(a)* Spatial correlation between cluster centroids reveals anticorrelation between DMNand DMN+, between DMNand FPN+, and between VIS+ and VIS-. *(b)* Spatial correlation between centroids calculated separately for rest and n-back reveal high correspondence, consistent with the identification of recurrent activity patterns common to both scans. *(c)* Cluster centroids computed by including equal amounts of rest and n-back task data as input to the clustering algorithm. Cluster names based on maximum cosine similarity were identical to the full sample centroids. *(d)* Spatial correlation between the 6 minute rest and the 6 minute n-back task cluster centroids and the full sample cluster centroids. Correlations > 0.99 were found only on the diagonal, suggesting 1-to-1 correspondence between the two centroid sets. The observed off-diagonal anticorrelations are consist with those observed in the full sample, as shown in panel *(a)*. *(e)* Group average state transition probabilities for rest (*right)* and n-back (*left)* using the 6 minute n-back task cluster partition reveals similar structure and high correlation with full sample state transition probabilities. *(f)* The *y*-axis shows the percentage of subjects missing at least one state in their time series for rest (*purple*) and for n-back (*yellow)*, for values of *k* on the *x*-axis ranging from 2 to 18. There existed at least one subject with missing states for all *k >* 5, supporting the choice of *k* = 5 for the main text.

**Fig. S3.**
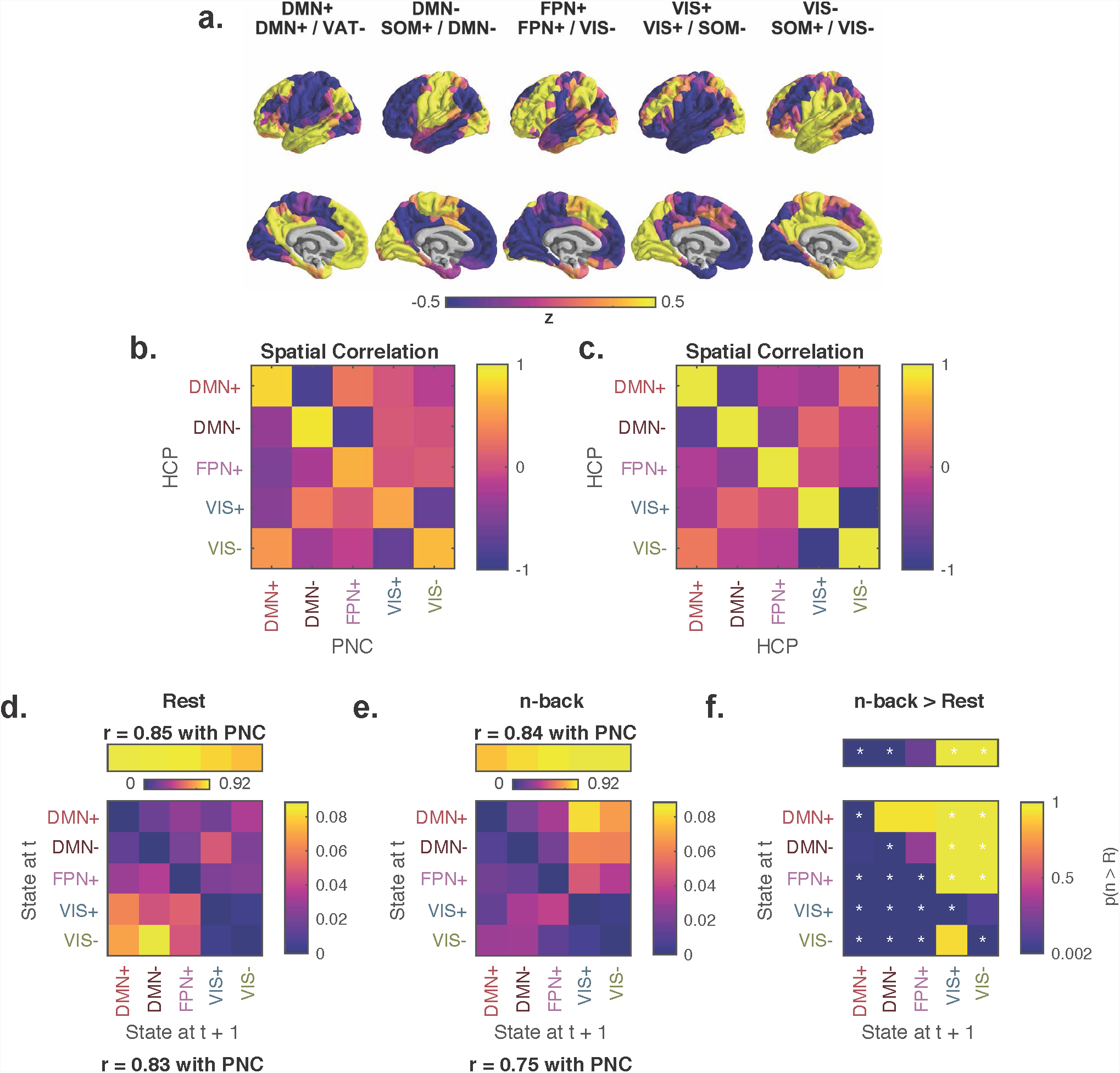
Brain states and dynamics in an independent sample with higher sampling rate and no global signal regression. *(a)* Cluster centroids for clustering of rest and n-back task BOLD data from the Human Connectome Project (HCP) with volumes acquired 4 times as frequently as the PNC and no global signal regression. *(b)* Spatial correlation between centroids for HCP and PNC data sets, is high along the diagonal, allowing for unambiguous matching of brain states between the two samples. *(c)* Spatial correlation between HCP centroids. DMN+ and DMN-, along with VIS+ and VIS-, exhibit strong anticorrelation. *(d-e)* HCP group average state transition probability matrices for rest *(d)* and n-back *(e)* scans. Off-diagonal elements of HCP rest and n-back transition matrices exhibit Pearson correlations of *r* = 0.83 and *r* = 0.75 with the PNC, respectively. HCP persistence probabilities are correlated with PNC persistence probabilities at *r* = 0.85 and *r* = 0.84 for rest and n-back, respectively. *(f)* Non-parametric permutation testing demonstrating differences between the rest and n-back group average transition probabilities and persistence probabilities. Extremes of the color axis indicate statistical significance, with larger values indicating higher transition probabilities in n-back relative to rest. *, Bonferroni-adjusted *p <* 0.05 or *p >* 0.95.

**Fig. S4.**
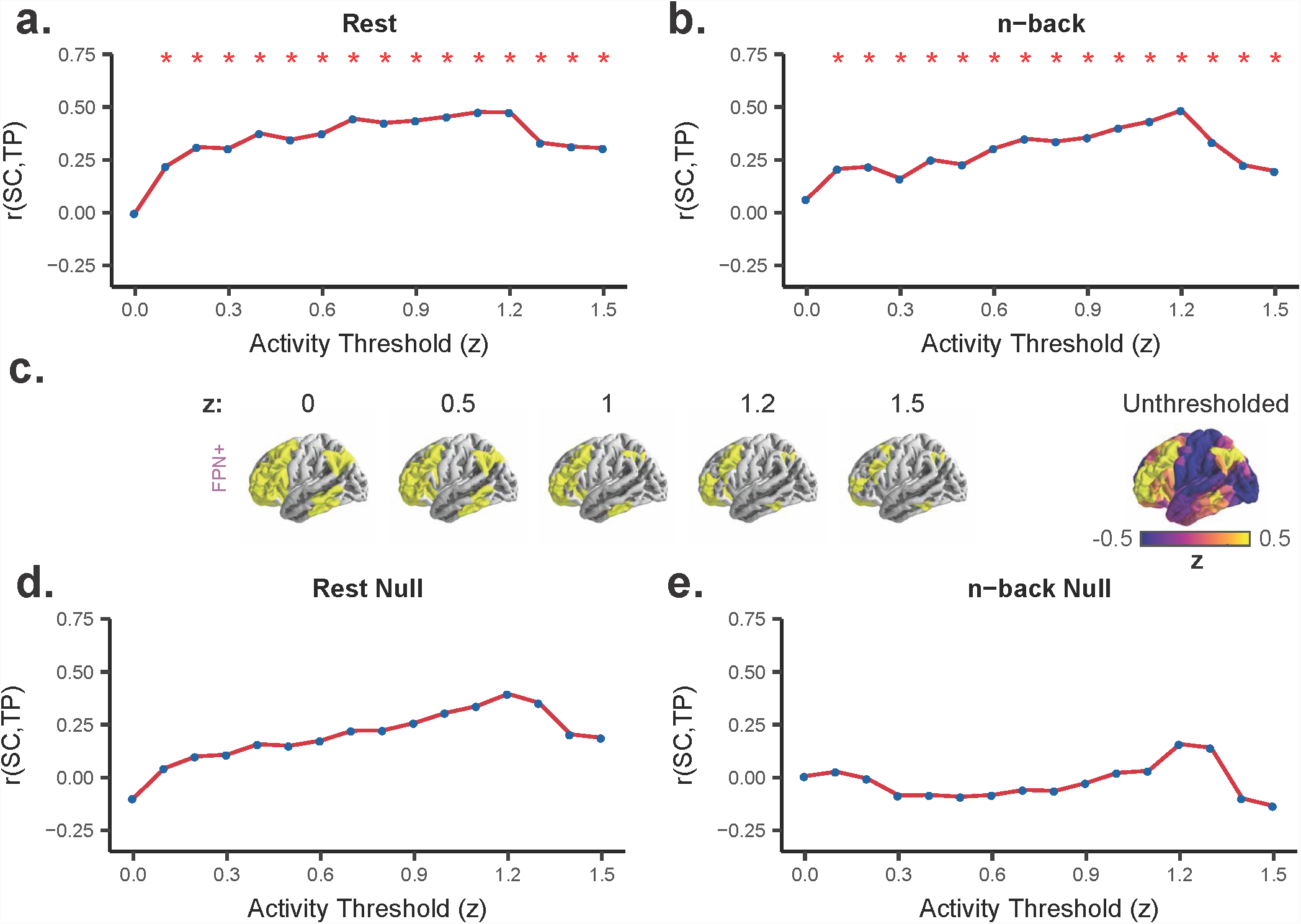
Structural connectivity predicts brain state transitions. *(a-b)* Correlation between group average inter-state structural connectivity (SC) and group average rest *(a)* or n-back *(b)* state transition probabilities for a range of threshold values defining active nodes. Red *, *p <* 10^-15^ for paired *t*-test comparing subject-level correlations between actual and subjectspecific topological null models^52^. *(c)* Example of procedure for determining active regions using a range of threshold values. Yellow regions exhibit activity greater than the *z*-score threshold for activity in the FPN+ cluster centroid. *(d-e)* Correlation between group average inter-state structural connectivity (SC) computed from subject-specific topological null models^52^ and group average rest *(d)* or n-back *(e)* state transition probabilities for a range of threshold values defining active nodes. *SC*, structural connectivity. *TP*, transition probability.

**Fig. S5.**
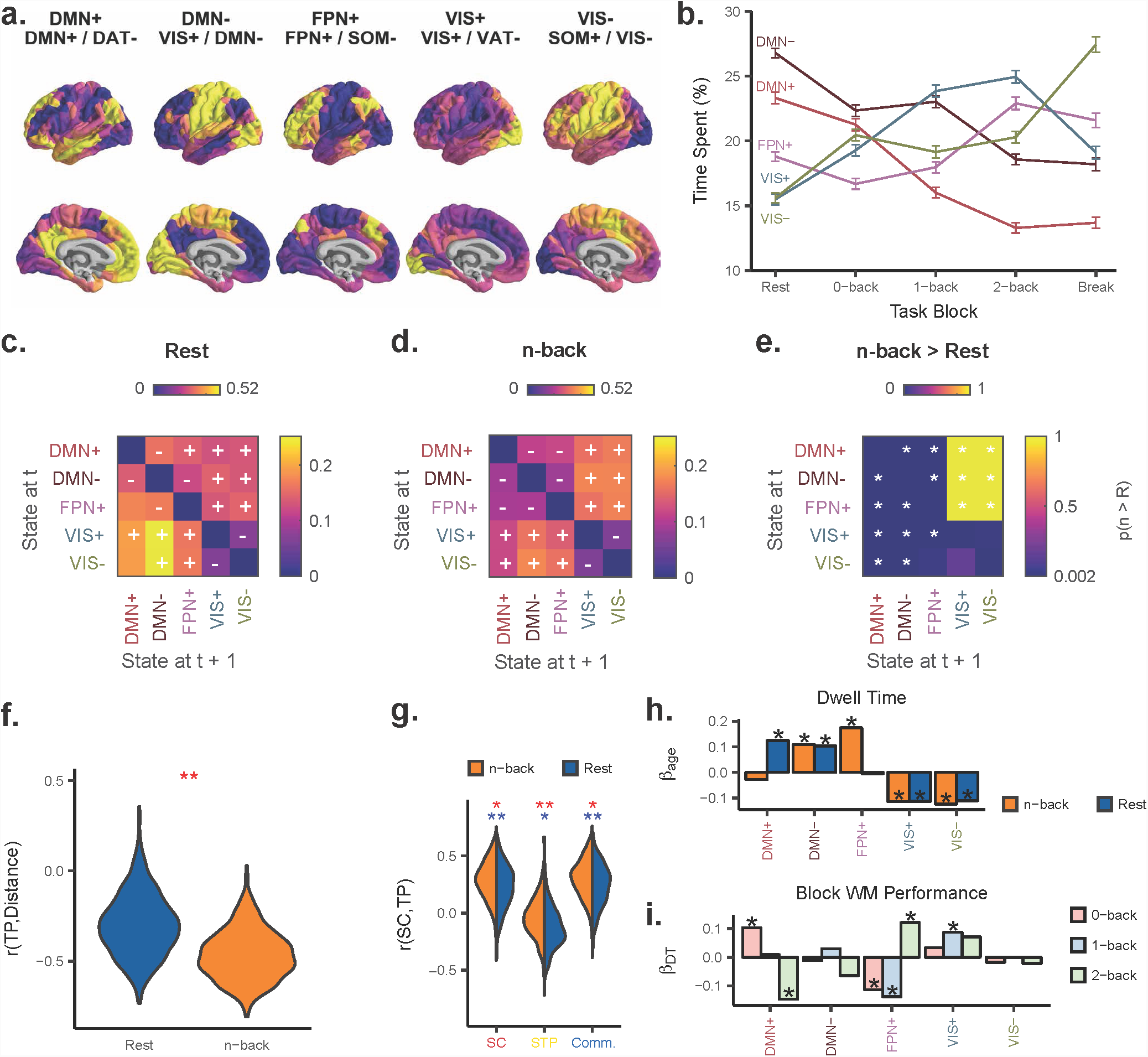
Key findings reproduced at *k* = 5 using cosine distance. *(a)* Cluster centroids at *k* = 5, generated from *k*-means clustering using cosine similarity as the distance function. *(b)* State dwell times change with increasing cognitive load. *(c-d)* Group average state transition probability matrices for rest (panel *(c)*) and n-back (panel *(d)*) scans. Overlayed + or indicates Bonferroni-adjusted *p <* 0.05 for state transitions occurring more or less, respectively, than expected under a null model. For visualization purposes, the state persistence probabilities are removed from the diagonal and depicted above the transition matrix. *(e)* Permutation testing to compare rest and n-back state transition probabilities. Values near 1 for a transition indicate n-back > rest, while values near 0 indicate rest > n-back. *, statistically significant after Bonferroni correction. *(f)* Correlation between transition probability and inter-state Euclidean distance is lower for n-back than for rest. **, *p <* 10^-15^. *(g)* Correlations between metrics of inter-state SC and state transition probabilities at rest and during n-back task using an activity threshold of *z* = 1.1. Blue *, *p <* 10^-6^ for *t*-test comparing the distribution to 0; blue **, *p <* 10^-15^ for *t*-test comparing the distribution to 0; red *, *p <* 10^-5^ for paired *t*-test comparing rest and n-back; red **, *p <* 10^-15^ for paired *t*-test comparing rest and n-back. *(h)* Standardized linear regression *β* weights for age as a predictor of dwell time in each state during rest and during n-back task performance, controlling for brain volume, handedness, head motion, and sex. *(i)* Standardized linear regression *β* weights for state-specific dwell time on working memory (WM) performance for each task block requiring an increasing WM load (0-back, 1-back, and 2-back), controlling for age, brain volume, handedness, head motion, and sex. For panels *(h-i)*, * indicates *p <.*05 after Bonferroni correction. *TP*, transition probability. *DT*, dwell time. *SC*, structural connectivity. *STP*, shortest topological path. *Comm.*, communicability. *(WM)*, working memory.

**Fig. S6.**
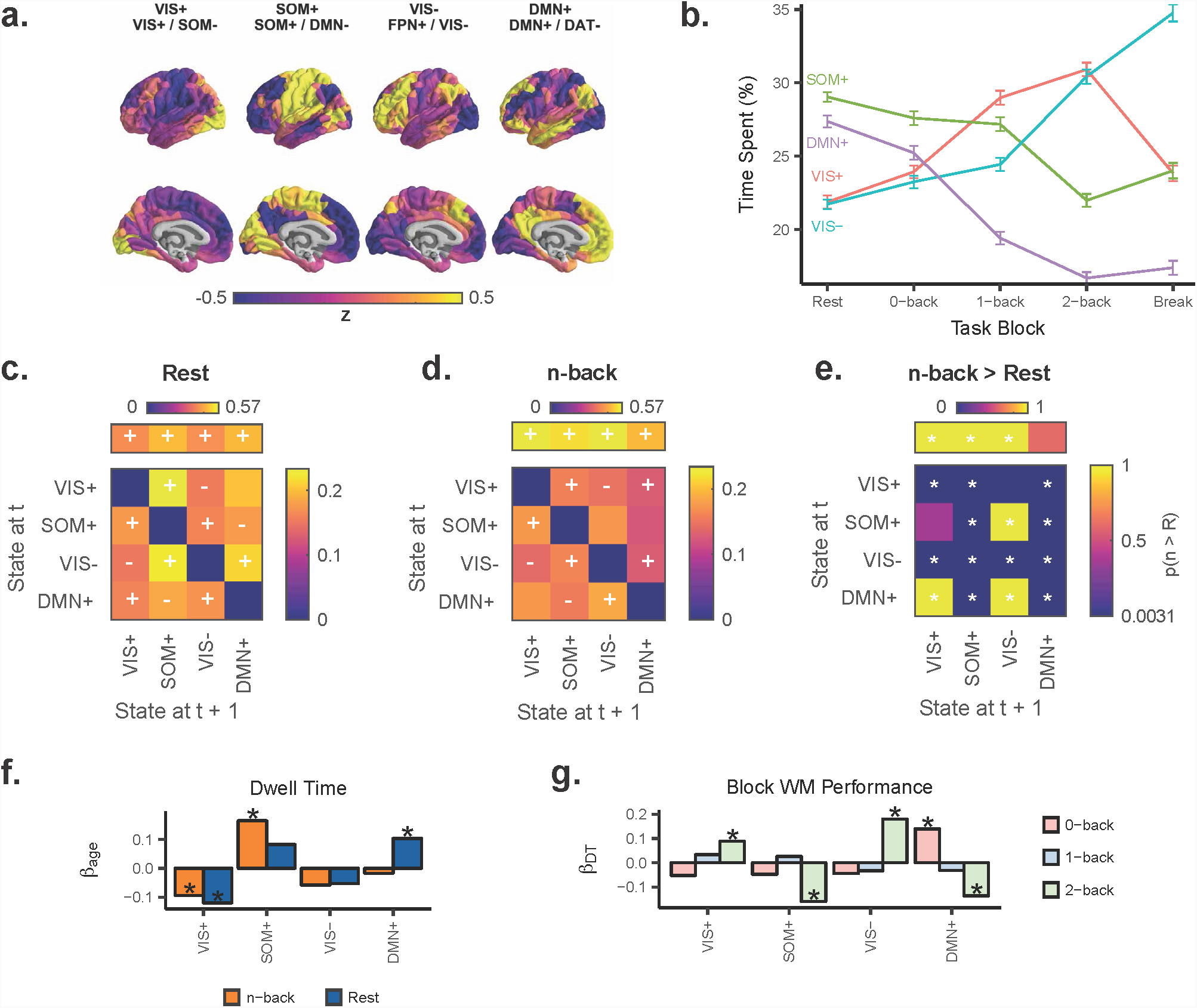
Key findings reproduced at *k* = 4. *(a)* Cluster centroids at *k* = 4, similar to *k* = 5 with FPN+ cluster folded into VIS-, among other changes. *(b)* State dwell times change with increasing cognitive load similarly compared to *k* = 5. *(c-d)* Group average state transition probability matrices for rest (panel *(c)*) and n-back (panel *(d)*) scans. Overlayed + or indicates Bonferroni-adjusted *p <* 0.05 for state transitions occurring more or less, respectively, than expected under an appropriate non-parametric null model. For visualization purposes, the state persistence probabilities are removed from the diagonal and depicted above the state transition matrix. *(e)* Permutation testing to compare rest and n-back state transition probabilities. Values near 1 for a state transition indicate n-back > rest, while values near 0 indicate rest > n-back. *, statistically significant after Bonferroni correction. *(f)* Standardized linear regression *β* weights for age as a predictor of dwell time in each state during rest and during n-back task performance, controlling for brain volume, handedness, head motion, and sex. *(g)* Standardized linear regression *β* weights for state-specific dwell time on working memory (WM) performance for each task block requiring an increasing WM load (0-back, 1-back, and 2-back), controlling for age, brain volume, handedness, head motion, and sex. For panels *(h-i)*, * indicates *p <.*05 after Bonferroni correction. *TP*, transition probability. *DT*, dwell time. *(WM)*, working memory.

**Fig. S7.**
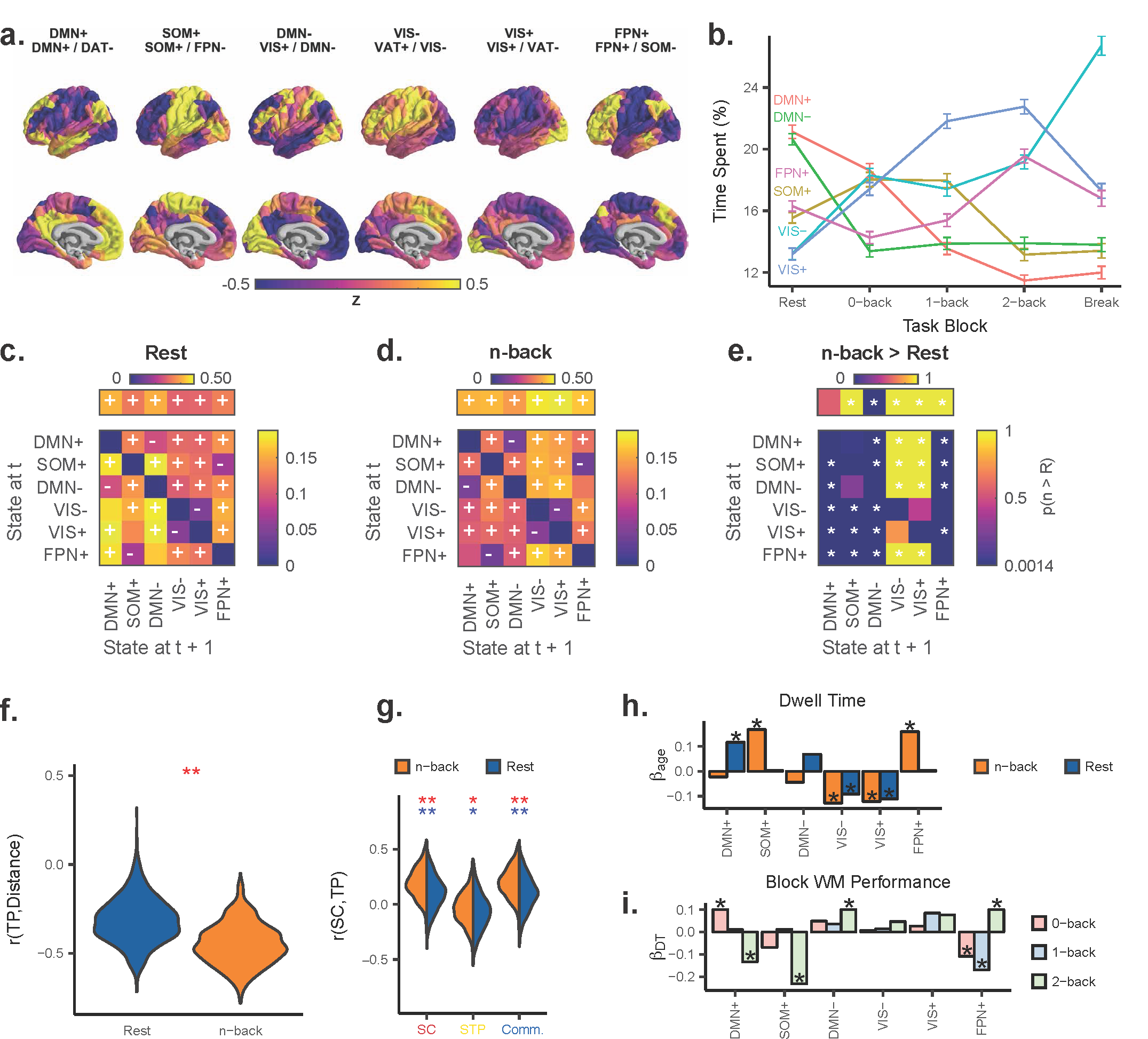
Key findings reproduced at *k* = 6. *(a)* Cluster centroids at *k* = 6, similar to *k* = 5 with the addition of a SOM+ cluster. *(b)* State dwell times change with increasing cognitive load similarly compared to *k* = 5. *(c-d)* Group average state transition probability matrices for rest (panel *(c)*) and n-back (panel *(d)*) scans. Overlayed + or indicates Bonferroni-adjusted *p <* 0.05 for state transitions occurring more or less, respectively, than expected under an appropriate non-parametric null model. For visualization purposes, the state persistence probabilities are removed from the diagonal and depicted above the state transition matrix. *(e)* Permutation testing to compare n-back and rest transition probabilities. Values near 1 for a state transition indicate n-back > rest, while values near 0 indicate rest > n-back. *, statistically significant after Bonferroni correction. *(f)* Correlation between state transition probability and inter-state Euclidean distance is lower for n-back than for rest. **, *p <* 10^-15^. *(g)* Correlations between metrics of inter-state SC and state transition probabilities at rest and during the n-back task using an activity threshold of *z* = 1. Blue *, *p <* 10^-6^ for *t*-test comparing the distribution to 0; blue **, *p <* 10^-15^ for *t*-test comparing the distribution to 0; red *, *p <* 10^-5^ for paired *t*-test comparing rest and n-back; red **, *p <* 10^-15^ for paired *t*-test comparing rest and n-back. *(h)* Standardized linear regression *β* weights for age as a predictor of dwell time in each state during rest and during n-back task performance, controlling for brain volume, handedness, head motion, and sex. *(i)* Standardized linear regression *β* weights for state-specific dwell time on working memory (WM) performance for each task block requiring an increasing WM load (0-back, 1-back, and 2-back), controlling for age, brain volume, handedness, head motion, and sex. For panels *(h-i)*, * indicates *p <.*05 after Bonferroni correction. *TP*, transition probability. *DT*, dwell time. *SC*, structural connectivity. *STP*, shortest topological path. *Comm.*, communicability. *(WM)*, working memory.

**Fig. S8.**
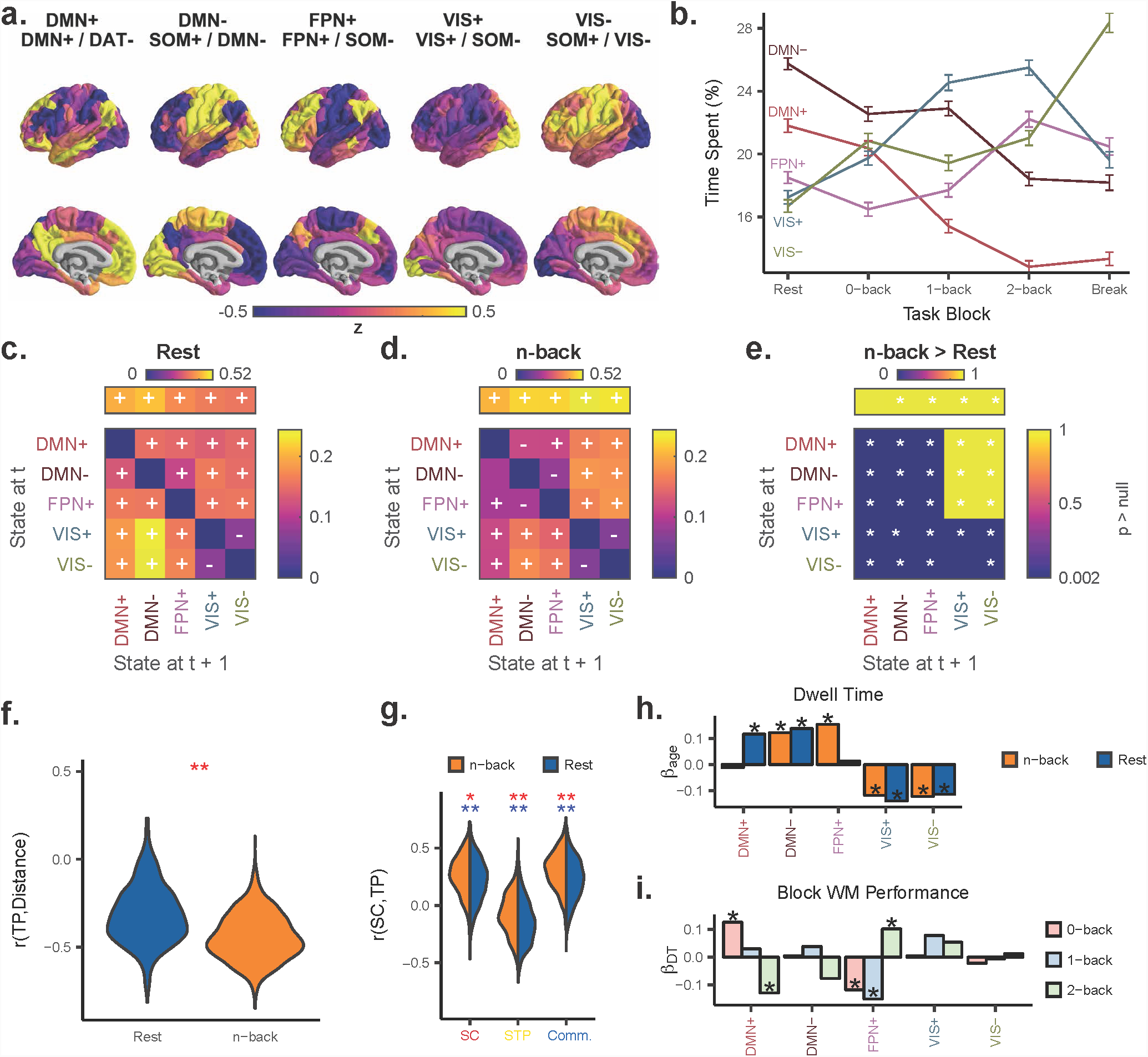
Key findings reproduced at *k* = 5 using the 234-node Lausanne parcellation. *(a)* Cluster centroids at *k* = 5 are similar to that of the 463-node Lausanne parcellation. *(b)* State dwell times change with increasing cognitive load similarly compared to the 463-node parcellation analysis. *(c-d)* Group average state transition probability matrices for rest (panel *(c)*) and n-back (panel *(d)*) scans. Overlayed + or indicates Bonferroni-adjusted *p <* 0.05 for state transitions occurring more or less, respectively, than expected under an appropriate non-parametric null model. For visualization purposes, the state persistence probabilities are removed from the diagonal and depicted above the state transition matrix. *(e)* Permutation testing to compare rest and n-back state transition probabilities. Values near 1 for a state transition indicate n-back > rest, while values near 0 indicate rest > n-back. *, statistically significant after Bonferroni correction. *(f)* Correlation between state transition probability and inter-state Euclidean distance is lower for n-back than for rest. **, *p <* 10^-15^. *(g)* Correlations between metrics of inter-state SC and state transition probabilities at rest and during the n-back task using an activity threshold of *z* = 1.1. Blue *, *p <* 10^-6^ for *t*-test comparing the distribution to 0; blue **, *p <* 10^-15^ for *t*-test comparing the distribution to 0; red *, *p <* 10^-5^ for paired *t*-test comparing rest and n-back; red **, *p <* 10^-15^ for paired *t*-test comparing rest and n-back. *(h)* Standardized linear regression *β* weights for age as a predictor of dwell time in each state during rest and during n-back task performance, controlling for brain volume, handedness, head motion, and sex. *(i)* Standardized linear regression *β* weights for state-specific dwell time on working memory (WM) performance for each task block requiring an increasing WM load (0-back, 1-back, and 2-back), controlling for age, brain volume, handedness, head motion, and sex. For *(h-i)*, * indicates *p <.*05 after Bonferroni correction. *TP*, transition probability. *DT*, dwell time. *SC*, structural connectivity. *STP*, shortest topological path. *Comm.*, communicability. *(WM)*, working memory.

**Fig. S9.**
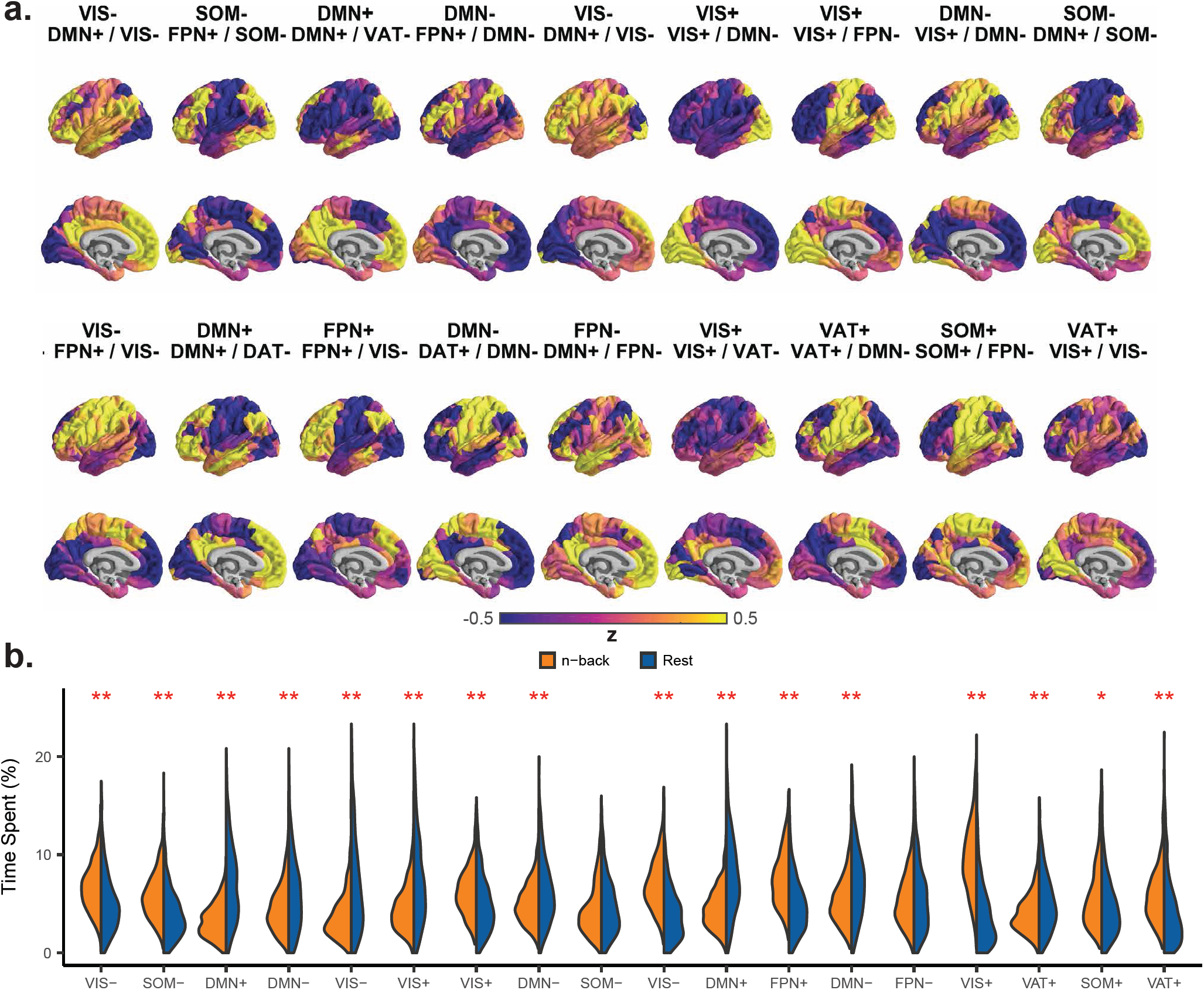
Brain states at *k* = 18. *(a)* Cluster centroids at *k* = 18 are reminiscent of brain states at *k* = 5 but with several additional combinations of resting state network activity patterns. *(b)* Nearly every state at *k* = 18 has a different dwell time for rest and n-back. *, *p <* 10^-4^, **, *p <* 10^-15^.

